# E-Selectin Orchestrates IL-1β–Dependent Neuroinflammation via NLRP3 in Vincristine-Induced Neuropathy

**DOI:** 10.64898/2026.03.10.710929

**Authors:** Hana Starobova, Ammar Alshammari, Nikhil Nageshwar Inturi, Nicolette Tay, Svetlana Shatunova, Ali Lam, Quan Nguyen, Marisol Mancilla Moreno, Diana Tavares-Ferreira, Federico Iseppon, Virginia Rodriguez-Menendez, Cristina Meregalli, Brittany Hill, Larissa Labzin, Simranpreet Kaur, Darren L. Brown, Guido Cavaletti, Theodore J. Price, Avril Robertson, Jennifer L. Stow, Ingrid G Winkler, Irina Vetter

## Abstract

Vincristine-induced peripheral neuropathy (VIPN) is a frequent and dose-limiting complication of cancer therapy, yet the upstream mechanisms coupling vascular activation to neuroinflammation remain poorly defined. Here we identify the vascular leukocyte adhesion molecule E-selectin (CD62E) as a critical orchestrator of vincristine-induced neuropathy. Systematic interrogation of endothelial-expressed leukocyte cell adhesion molecules (CAMs) in a murine model of VIPN revealed that blockade of E-selectin, but not ICAM-1, PECAM-1 or P-selectin, completely prevented mechanical hypersensitivity and markedly reduced F4/80 immune cell accumulation in dorsal root ganglia (DRG) and peripheral nerves. Genetic deletion of E-selectin (Sele) conferred similar VIPN protection, despite the absence of structural intraepidermal or myelinated fibres loss, indicating a predominantly functional neuroimmune pathology. Spatial transcriptomics demonstrated that vincristine induces a conserved stress and neuroinflammation-associated transcriptional programme in DRG, with immune and stromal populations acting as dominant signalling hubs. Genetic or pharmacological perturbation of E-selectin did not abolish injury-associated pathways but redistributed cell–cell communication networks, reducing immune-cell dominance and reshaping interferon and metabolic signalling. Mechanistically, E-selectin exerted non-canonical effects beyond endothelial adhesion. Local recombinant E-selectin administration was sufficient to induce phagocyte-dependent mechanical hypersensitivity that was abolished in Fut4/7- (functional receptor) deficient mice. In macrophages, E-selectin adhesion mediated enhanced vincristine-driven NF-κB activation, NLRP3 inflammasome assembly and IL-1β release. Together, these findings position E-selectin as an upstream regulator of IL-1β– dependent neuroinflammation in VIPN and identify selective targeting of E-selectin– mediated immune-neuron interactions as a potential preventative therapeutic strategy for chemotherapy-induced neuropathy.

## Introduction

Vincristine is a chemotherapeutic agent widely used in the treatment of paediatric and adult malignancies, including acute lymphoblastic leukaemia, Hodgkin’s lymphoma, and neuroblastoma^1–3^. However, its clinical utility is frequently constrained by vincristine-induced peripheral neuropathy (VIPN), a dose-limiting and often irreversible adverse condition that affects up to 90% of paediatric patients and a substantial proportion of adults^4^. VIPN presents with a spectrum of sensory, motor, and autonomic disturbances that significantly impair quality of life and long-term functional outcomes in cancer survivors^5,6^.

Although the underlying mechanisms of VIPN remain poorly understood, it is thought to involve microtubule disruption, impaired axonal transport, axonal degeneration, and mitochondrial dysfunction (reviewed in^6^). Recent studies have implicated neuroinflammation as a central driver of VIPN pathology^7–9^. Specifically, monocyte infiltration into dorsal root ganglia (DRG) and peripheral nerves initiates a cascade of immune activation, including the assembly of the NLRP3 inflammasome and release of pro-inflammatory cytokines such as interleukin-1β (IL-1β), a critical mediator of inflammatory pain^8,10–12^. Relevant to the clinic, inhibition of the NLRP3 inflammasome using MCC950, a selective NLRP3 inhibitor^13^, alleviates vincristine-induced sensory and motor neuropathy in ALL-19 tumour-bearing NSG mice treated with a combination of vincristine, L-asparaginase, and dexamethasone^14^. However, while NLRP3-dependent inflammation has emerged as a key effector of vincristine neurotoxicity, the upstream signals that govern immune cell recruitment to peripheral nerves remain poorly defined.

Vincristine is known to upregulate the expression of intercellular adhesion molecule-1 (ICAM-1) and vascular cell adhesion molecule-1 (VCAM-1)^7^, and it is plausible that enhanced leukocyte adhesive properties of endothelial cells help drive the infiltration of immune cells into neuronal tissue. However, the functional contribution of different families of vascular cell adhesion molecules (CAMs), such as the integrin ligands and selectins, to the development of vincristine-induced neuropathy has not been systematically assessed. To address this gap, we investigated the contributions of VCAM-1 (CD106,) ICAM-1 (CD54), PECAM-1 (CD31), P-selectin (CD62P) and E-selectin (CD62E) to vincristine-induced mechanical hypersensitivity and neuronal immune cell infiltration.

Using a murine model of VIPN, we identify E-selectin as a critical driver of VIPN. Pre-treatment with an E-selectin–blocking antibody provided complete protection against vincristine-induced mechanical hypersensitivity, highlighting its essential role in neuropathy development. Moreover, E-selectin inhibition or genetic deletion conferred neuroprotection on transcriptional level in the DRG. Importantly, our findings also uncover a non-canonical signalling role of E-selectin in VIPN, where, in addition to immune cell infiltration, it promotes neuroinflammation by enhancing NF-κB activation and amplifying IL-1β release through the NLRP3 inflammasome.

## Materials and methods

For additional information on materials and methods see Supplementary materials.

### Animals, in vivo experiments and ethical approvals

Behavioural and in vivo experiments were performed using 8–10-week-old male C57BL/6J, E-selectin-deficient (Sele^−/−^), fucosyltransferase 4/7 double-knockout (Fut4^−/−^Fut7^−/−^), and ASC-citrine reporter mice. The Sele^−/−^ and Fut4^−/−^Fut7^−/−^ lines were maintained on a C57BL/6 background and have previously been reported to have been backcrossed onto this background for at least 10 generations^15^. The C57BL/6 background of Fut4^−/−^Fut7^−/−^ mice has also been independently reported^16^. ASC-citrine reporter mice were generated and characterised as previously described^17^. Age-matched C57BL/6J mice, rather than littermate wild-type controls, were used as controls for knockout experiments. Only male mice were used as no sex differences in VIPN were observed previously^18^. Mice were group-housed (3– 5 per cage) under a 12 h light/dark cycle with ad libitum food and water. All procedures were approved by the University of Queensland Animal Ethics Committee and conducted in accordance with the Animal Care and Protection Regulation (Qld, 2012), the Australian Code for the Care and Use of Animals for Scientific Purposes (2013), and IASP guidelines. Animals were acclimatised prior to testing, randomly assigned to treatment groups, and behavioural assessments were performed by blinded observers. Vincristine sulphate (Pfizer) was diluted in sterile phosphate-buffered saline (PBS) and administered intraperitoneally (i.p.) at 0.5 mg/kg (10 μl/g), as previously described^8^. Blocking antibodies against E-selectin, VCAM-1, ICAM-1, PECAM-1 (Bio X Cell), or P-selectin (BioLegend), or matching isotype controls (10 mg/kg; 10 μl/g) as well as anakinra (100 mg/kg, Sobi) were administered prior to vincristine administration. For intraplantar injections, recombinant mouse E-selectin (600 ng/injection; R&D Systems), human IgG isotype control (600 ng/injection; Invitrogen) or bone marrow-derived macrophages (BMDMs) were diluted in sterile PBS and injected intraplantar (i.pl.; 1000 cells/20 μl/paw) into the right hind paw of C57BL/6J or *Fut4/7 /* mice under 2% isoflurane anaesthesia. Mechanical and thermal sensitivity of the hind paws was assessed using electronic von Frey and MouseMet thermal apparatuses (MouseMet; Topcat Metrology) as previously described^18^. To deplete phagocytic cells, mice received intraperitoneal (i.p.) injections of liposome-encapsulated clodronate (10 μl/g of 5 mg/mL; Liposoma) or control liposome-encapsulated PBS as described previously^8^.

### Animal selection, allocation and sample size

Male mice aged 8–10 weeks with the genotype required for each experiment were eligible for inclusion. Before allocation, mice were assessed for general health and for conditions that could interfere with behavioural testing or tissue analysis, including visible paw injury, impaired mobility unrelated to treatment, or illness. An independent researcher randomly assigned mice to treatment groups and retained the allocation key. Investigators conducting behavioural assessments, histological quantification and other analyses remained blinded to group identity until analyses were complete. All animals assigned to experimental groups received their allocated treatment and were retained in the study; no animals were excluded after allocation.

Behavioural assessments, histological quantification, and other analyses were performed by investigators blinded to treatment allocation. The mouse was the experimental unit for in vivo outcomes. Repeated paw withdrawal measurements and multiple tissue sections from each mouse were averaged to generate one value per mouse for each endpoint and time point; these measurements were not treated as independent biological replicates.

Group sizes for behavioural experiments were selected based on 80% power to detect a 60% difference in the primary outcome relative to baseline, assuming 20% interindividual variability and P < 0.05 and effect sizes and variability observed in our previously published vincristine model ^8^, together with the number of treatment groups and animal-use considerations. Sample sizes for histological and cellular assays were selected based on prior experience with these endpoints and are stated separately in each figure legend. The spatial transcriptomic experiment included three independent mice per condition and was intended to identify candidate changes for mechanistic interpretation; individual spatial spots were not considered biological replicates.

### Quantification of cytokine release and ASC speck formation in BMDMs

Murine (C57BL6/J or ASC-citrine) BMDMs were isolated and cultured as previously described^19,20^. Cytokine and chemokine release was quantified in unadhered BMDMs or following adhesion of BMDMs to plates coated with recombinant mouse E-selectin (5 μg/mL; 575-ES, R&D Systems), recombinant mouse P-selectin (5 μg/mL; R&D Systems), recombinant human IgG isotype control (5 μg/mL; Invitrogen), or Tris-buffered saline^21^ as a control by LEGENDplex^TM^ Mouse Macrophage/Microglia Panel (BioLegend) according to the manufacturer’s protocol. Readouts from samples were obtained using a flow cytometer (Beckman Coulter CytoFLEX-S). Data were analysed using the Data Analysis Software Suite by BioLegend. The concentration of interleukin-1 beta (IL-1β) from ASC-Citrine reporter BMDMs was quantified using ELISA kit (R&D Systems) according to the manufacturer’s protocol. For ASC speck quantification, live-cell imaging was performed every 10 min for up to 220 min using a Nikon Ti-E inverted microscope with a Hamamatsu Flash 4.0 sCMOS camera.

### Quantification of DRG neuron and BMDM adhesion to E-selectin

Dorsal root ganglia (DRGs) were isolated from 8-week-old male C57BL/6 mice as previously described ^19,20,22,23^. Dissociated DRG neurons or BMDMs were seeded into 96-well plates pre-coated with recombinant mouse E-selectin, isotype control, or Tris-buffered saline (5 μg/mL; 24 h). Cell suspensions were added and incubated at 4°C for 1 h to allow adhesion. Non-adherent cells were removed by washing three times with fresh DMEM-high glucose (40 μL/well). Adherent cells were counted under a light microscope by a blinded observer.

### Nf-κB p65 Transcription Factor Assay

NF-κB activation was assessed using the NFκB p65 Transcription Factor Assay Kit (Abcam) following the manufacturer’s instructions. Nuclear proteins were extracted from BMDMs 30 min post-treatment using the Abcam Nuclear Extraction Kit. Protein concentrations were measured via Nanodrop, and 20 µL of nuclear extract per sample was analysed in duplicate. Activated NF-κB p65 binding to DNA was detected using a specific antibody, HRP-conjugated secondary antibody, and a colorimetric readout at 450 nm. Absorbance was measured using a Tecan microplate reader, and data were analysed to compare NF-κB activity between conditions.

### Immunohistochemistry and quantification of Intraepidermal Nerve Fibre (IENF) density

F4/80 staining was performed using a previously optimised and validated immunohistochemical protocol for mouse DRG ^8,24,25^. Sections were stained with rat anti-mouse F4/80, developed using ABC/DAB, counterstained with haematoxylin, scanned, and quantified by a blinded observer using Visiopharm. Representative images are provided in Supplementary Figure S1. Briefly, dorsal root ganglia and sciatic nerves for immunohistochemistry were collected 25 h post vincristine or vehicle injection, fixed in 10% neutral-buffered formalin (NBF), and embedded in paraffin as previously described^25^. Briefly, Paraffin sections (5 μm) were deparaffinised, rehydrated, and antigen-retrieved with Proteinase K. After blocking with Background Sniper (BioCare Medical), sections were incubated with rat anti-mouse F4/80 (1:350; Novus Bio) for 2 h, followed by biotinylated secondary antibody, ABC amplification (Agilent Dako), and DAB detection. Nuclei were counterstained with haematoxylin. Isotype control (rat IgG2b; BioLegend) was used to confirm specificity. Slides were scanned (Olympus) and F4/80^+^ area quantified using Visiopharm by a blinded observer to assess F4/80^+^ cell infiltration across treatment groups. F4/80-positive area was quantified as the percentage of the analysed tissue area occupied by thresholded DAB-positive F4/80 immunoreactivity.

For quantification of intraepidermal nerve fibre (IENF) density, vincristine (0.5 mg.kg. i.p.; Pfizer) was injected 5 times a week for 4 weeks. 1 week after the final vincristine injection (cumulative dose: 10 mg/kg), plantar hind paw skin was collected, postfixed in 4% paraformaldehyde for 24 h at 4 °C, cryoprotected in 30% sucrose o/n, embedded in OCT (TissueTEK) and snap frozen using dry ice ^26^. Tissue was sectioned into three non-consecutive 50 µm sections per animal (n = 4 animals per group) and stained using rabbit anti-PGP9.5 primary antibody (1:500; Dako, Z5116) followed by Alexa Fluor 555–goat anti-rabbit secondary antibody (1:1000; Invitrogen, A-21428) ^26^. Sections were coverslipped with DAPI-containing mounting media (Fluoroshield, Sigma Aldrich, F6057). Images were acquired using a spinning-disc confocal microscope (Nikon Ti2/Andor Dragonfly). Image analysis was performed in FIJI (ImageJ). Intraepidermal nerve fibre density was expressed as the percentage of PGP9.5-positive area relative to total epidermal area.

### Spatial transcriptomics

Eight-week-old male C57BL/6J mice were treated with either saline or vincristine (0.5 mg/kg, i.p.) once. Additional groups included C57BL/6J mice pre-treated with an E-selectin-blocking antibody (9A9, BioX Cell, 200 µg/mouse, i.p. at two time points (-24h ,-1h) prior to vincristine), and E-selectin knockout (*Sele /* ) mice treated once with vincristine (0.5 mg/kg, i.p.). Dorsal root ganglia (DRG) were collected 24h post vincristine treatment under RNase-free conditions and immediately embedded in chilled OCT compound (Tissue-Tek, Sakura, Japan), then flash-frozen in pre-cooled isopentane to preserve tissue morphology. The study included DRGs from lumbar level L3-L5 (both sides) from 3 mice per treatment condition. A total of 15 DRGs per condition were embedded in one OCT block, positioned on a uniform embedding plane for consistent sectioning. Blocks were stored at – 80 °C until further processing. Cryosections were dried at 37 °C for 1 min, fixed in 100% methanol at –20 °C for 30 min, and stained with Mayer’s haematoxylin (5 min) and eosin (2 min). Slides were mounted in 85% glycerol for imaging on a Zeiss Axio Z1 slide scanner. Library preparation followed the Visium Spatial Gene Expression User Guide (CG000239 Rev C, 10x Genomics). Sequencing was performed using an Illumina NextSeq 500 with the following read structure: Read1 – 28 bp, Index1 – 10 bp, Index2 – 10 bp, Read2 – 120 bp. Raw BCL files were processed using bcl2fastq (v2.7.0), and reads were aligned to the mouse reference genome, quantified and assigned to spatial barcodes using Space Ranger (v1.3.0, 10x Genomics).

### Pseudo-bulk generation from Visium data and differential expression analysis

To ensure that statistical inference was performed at the level of independent biological replicates rather than individual Visium spots, barcodes originating from the same animal were aggregated to generate one pseudobulk count matrix per animal (**n = 3 biological replicates per condition**). Individual Visium spots were not treated as independent biological replicates. Differential gene expression analysis was performed using DESeq2 on raw gene counts ^27^. Size factors were estimated with the median ratio method (each gene count divided by a barcode specific factor derived from the geometric mean across samples), after which a batch term was included in the design formula to correct for sequencing batch effects (design = ∼ batch + condition). DESeq2 fits a negative binomial generalized linear model and uses the Wald test followed by Benjamini-Hochberg correction to identify differentially expressed genes. Log fold changes were shrunk using the “ashr” method^28^, improving effect size estimation for low count genes. Genes meeting adjusted P< 0.05 and log FC ≥ 0.585 (FC of 1.5) were reported as differentially expressed.

### Spatial Transcriptomics Deconvolution with SPOTlight

Cell-type composition of Visium spots was inferred using seeded non-negative matrix factorization (NMF) regression-based deconvolution implemented in **SPOTlight** (v1.8.0) ^29^. A harmonized cross-species single-cell RNA-seq dataset containing previously annotated neuronal and non-neuronal DRG populations was used as the reference^30^. Gene expression and metadata from the spatial transcriptomics dataset were used to construct a SpatialExperiment object, and marker-gene profiles from the reference clusters were used to generate cell-type-specific topic profiles. These profiles were projected onto individual Visium spots to estimate the relative contribution of each cell population. Thus, cell-type assignments represent reference-guided estimates of cellular composition rather than single-cell measurements and were used for spatial visualization and downstream cell-type-resolved analyses.

### Cell-cell interactions

Cell–cell communication analysis was performed using CellChat ^31^. The merged spatial Seurat object was split by condition to generate separate CellChat objects, and ligand– receptor interactions were inferred using the mouse secreted signalling database. Communication probabilities were calculated using the tri-mean method, with interactions involving fewer than 10 cells excluded. Pathway-level communication networks were inferred and aggregated to compare signalling strength and interaction numbers across cell types. For selected pathways, CellChat-derived communication probabilities and network centrality metrics were used to identify predominant sender, receiver, mediator and influencer populations and contributing ligand–receptor interactions. As CellChat communication probabilities are model-derived measures integrating ligand and receptor expression across interacting cell populations, they represent predicted relative communication strength rather than conventional gene-expression values and were therefore not interpreted or presented as gene-expression fold changes between conditions.

### Morphometrical evaluation of nerves

C57BL/6J or *Sele*^−/−^ animals were treated with vincristine (0.5 mg/kg; i.p.; Pfizer) 5 times a week for 4 weeks as described previously ^8^. In week 5, animals were culled using CO_2_ euthanasia and caudal nerves were collected from 3 animals/groups and fixed by immersion with 3% Glutaraldehyde in phosphate buffer solution (0.12M), post-fixed in 1% OsO_4_, and embedded in epoxy resin to be ready for morphometric examination, according to previously reported protocols ^32,33^. 1.5-µm–thick semi-thin sections of the resin embedded nerves out of 3 animals/group were then cut and stained with methylene blue. Images were acquired with a Nexscope Ne920 AUTO light microscope using a 60X magnification objective. (TiEsseLab Srl, Milano, Italy). The acquired images were then analysed using Image Pro-Plus software, a computer-assisted image analyser (Media Cybernetics). The density (fibres/mm^2^) and the axonal diameters of myelinated fibres (used to calculate the frequency distribution histograms) were measured in randomly selected sections again according to previously reported methods ^32,33^.

### Data and statistical analyses

The mouse was the independent biological unit for behavioural, histological and nerve-morphometry analyses. For behavioural outcomes, three paw withdrawal measurements collected at each time point were averaged to obtain one value per mouse per time point. For histological outcomes, measurements from multiple sections or images were averaged to obtain one value per mouse for each tissue and endpoint. Sections and images were not treated as independent biological replicates. For spatial transcriptomics, the mouse was the biological replicate (n = 3 per condition); counts from spots assigned to the same mouse were aggregated before pseudobulk differential-expression analysis. For BMDM experiments, an independent donor mouse/BMDM preparation was the biological replicate; replicate wells and imaging fields from that preparation were averaged within condition before statistical analysis. Duplicate wells in the NF-κB assay were technical replicates and were averaged to yield one value per independent BMDM preparation.

Non-transcriptomic analyses were performed in GraphPad Prism version 10. Longitudinal behavioural data were analysed by two-way repeated-measures ANOVA, with treatment group as the between-mouse factor and time as the within-mouse factor; the treatment × time interaction was included. Prespecified comparisons between treatment groups at individual time points were adjusted using Šídák’s multiple-comparisons test. Single-time-point outcomes were analysed using one-way ANOVA followed by the stated multiple-comparisons test, or a two-sided unpaired t-test for two groups, as specified for each figure. Dunnett’s test was used when several conditions were compared with one designated control; Šídák’s test was used for the specified pairwise comparisons. Tests were two-sided, and adjusted P < 0.05 was considered significant. Data are presented as mean ± SEM; n denotes independent biological replicates, with the exact n reported for each analysis.

## Results

### Inhibition of E-selectin and VCAM-1 mitigates vincristine-induced hypersensitivity and F4/80^+^ immune cell accumulation in neuronal tissue

Several studies have reported that the development of vincristine-induced peripheral neuropathy (VIPN) is associated with an increased presence of F4/80 cells in the dorsal root ganglia (DRGs) and peripheral nerves^7,8,34,35^. This accumulation is likely driven by the infiltration of CX3CR1 monocytes into peripheral nerves^7^. Vincristine appears to facilitate this infiltration by enhancing the adhesive properties of endothelial cells, including through upregulating the expression of intercellular adhesion molecule-1 (ICAM-1) and vascular cell adhesion molecule-1 (VCAM-1)^7^. However, the role of other endothelial adhesion molecules, particularly selectins in facilitating monocyte infiltration or expansion in peripheral nerves remains unclear, as is the contribution of these processes to the development of VIPN.

To investigate these questions, we administered blocking antibodies against E-selectin (E-sel Ab), P-selectin (P-sel Ab), VCAM-1 (VCAM1 Ab), ICAM-1 (ICAM1 Ab), and PECAM-1 (PECAM1 Ab) to vincristine-treated C57BL6/J mice and quantified both the number of F4/80-positive cells in DRGs and sciatic nerve, as well as impact on mechanical hypersensitivity that arises as a result of vincristine treatment.

Consistent with previous studies reporting up-regulation of VCAM-1 following vincristine treatment, inhibition of VCAM-1 significantly alleviated vincristine-induced mechanical hypersensitivity at 5 h and 24 h post vincristine administration (**Fig. 1A; Supplementary Table 1)** and also significantly reduced F4/80 staining, indicative of monocyte/macrophage infiltration, in both DRGs and sciatic nerves (**Fig. 1B, Fig. S1, Supplementary Table 2**). In contrast, inhibition of ICAM-1 did not significantly alleviate vincristine-induced hypersensitivity (**Fig. 1C, Supplementary Table 1**), nor prevented infiltration of F4/80^+^ cells in DRGs or sciatic nerves (**Fig. 1D**; **Fig. S1** and **Supplementary Table 2)**. PECAM-1 inhibition only partially alleviated vincristine-induced hypersensitivity (**Fig. 1E, Supplementary Table 1**) and did not prevent infiltration of F4/80^+^ cells in DRGs or sciatic nerves (**Fig 1F, Fig. S1** and **Supplementary Table 2)**. Similar results were also observed following treatment with a P-selectin blocking antibody, which partially improved the vincristine-induced decrease in paw withdrawal threshold at 5 hours but not at 24 h compared to the IgG2a (**Fig. 1G; Supplementary Table 1**), and had no significant effect on the number of F4/80 cells in the DRGs or sciatic nerve (**Fig. 1H; Fig. S1** and **Supplementary Table 2**).

**Fig. 1.**
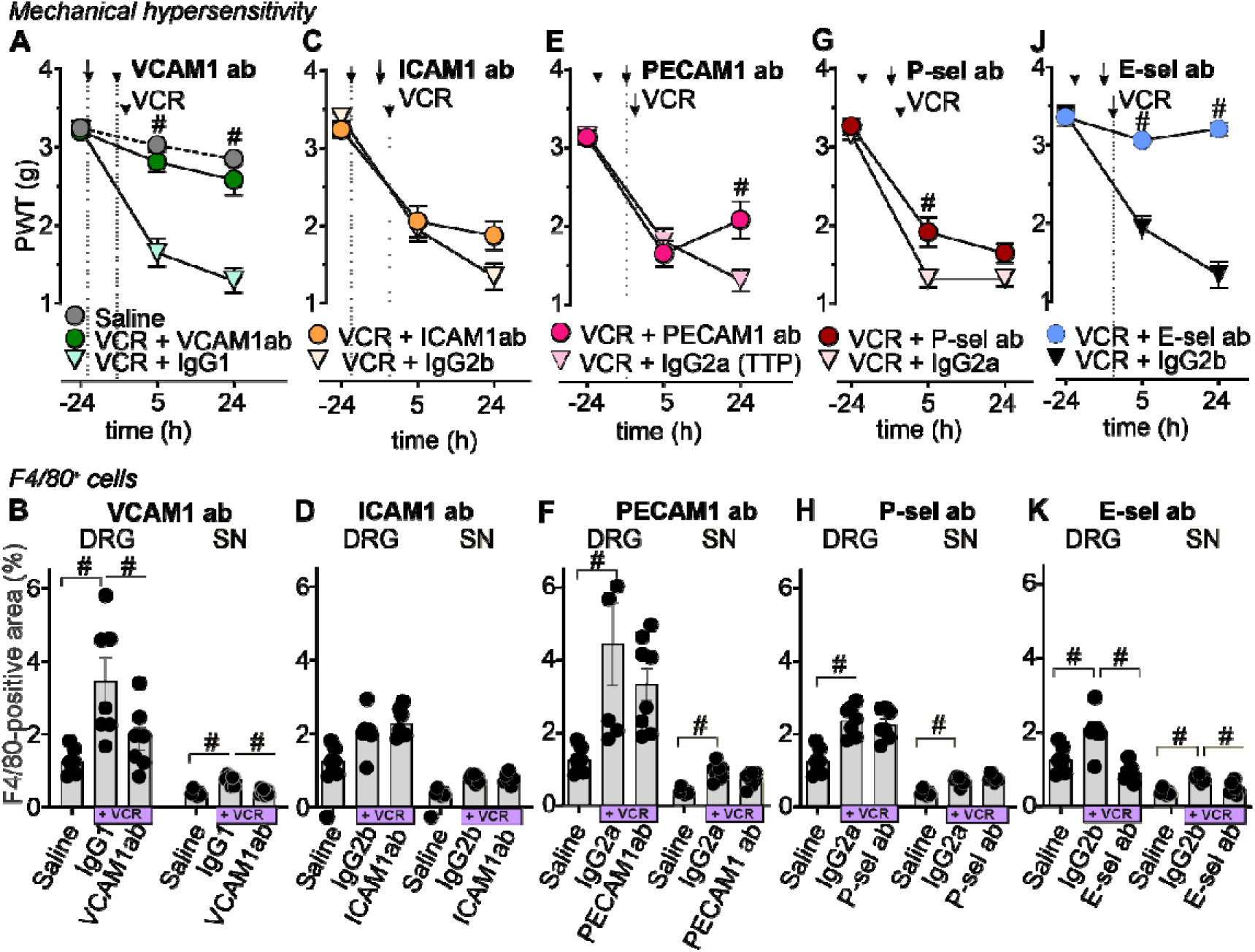
Inhibition of E-selectin and VCAM-1, but not other adhesion molecules, attenuates vincristine-induced mechanical hypersensitivity and increased F4/80-positive area in the dorsal root ganglia and sciatic nerves. Effects of blocking antibodies against VCAM-1 (VCAM1ab; **A,B**), ICAM-1 (ICAM1ab; **C,D**), PECAM-1 (**E,F**), P-selectin (P-sel ab; **G,H**) and E-selectin (E-sel ab; **J,K**) on vincristine-induced mechanical hypersensitivity (A,C,E,G,J; PWT, paw withdrawal threshold) and F4/80-positive area in the dorsal root ganglia (DRG) and sciatic nerve (SN) (B,D,F,H,K). Black arrows indicate i.p. administration of blocking antibodies (10 mg/kg), isotype control antibodies (IgG2b or IgG1; 10 mg/kg) and vincristine (0.5 mg/kg). VCAM-1 and E-selectin blockade attenuated vincristine-induced mechanical hypersensitivity (**A,J**) and the associated increase in F4/80-positive area in the DRG and SN (**B,K**), whereas PECAM-1 and P-selectin blockade was partially effective (**E–H**) and ICAM-1 blockade had no effect (**C,D**). F4/80-positive area was quantified as the percentage of analysed tissue area occupied by thresholded DAB-positive F4/80 immunoreactivity and does not represent the percentage or number of F4/80-positive cells. Data are presented as mean ± SEM (n = 5–7 per group). Statistical significance (#P < 0.05) was determined using repeated-measures two-way ANOVA or one-way ANOVA with Šídák’s multiple-comparisons test.

Interestingly, inhibition of E-selectin prior to vincristine administration significantly (compared to the IgG2b) and completely prevented development of vincristine-induced mechanical hypersensitivity (**Fig. 1J; Supplementary Table 1),** and also significantly reduced the number of F4/80 cells in both the DRGs and sciatic nerves (**Fig. 1K; Fig. S1** and **Supplementary Table 2**).

### Vincristine-induced mechanical hypersensitivity is attenuated in Sele□/□mice

Although E-selectin has been highlighted by a number of studies as a putative biomarker for pain (e.g. Study of Intravenous GMI-1070 in Adults with Sickle Cell Disease, NCT00911495) it has not previously been implicated as an analgesic target in chemotherapy-induced neuropathy. We thus sought to further investigate the molecular mechanisms contributing to anti-allodynic effects of E-selectin inhibition, as well as the potential of E-selectin-targeting therapeutics for treatment or prevention of vincristine-induced neuropathy.

We first sought to confirm that the phenotype observed following inhibition of E-selectin signalling with blocking antibodies – decreased immune cell infiltration in neuronal tissues, and decreased vincristine-induced mechanical hypersensitivity – also occurs in E-selectin knockout (*Sele*^−/−^) mice. Indeed, *Sele /* mice were fully protected from vincristine-induced mechanical hypersensitivity, whereas C57BL6/J (wt) mice exhibited a sustained decline in paw withdrawal thresholds (PWTs) (**Fig. 2A**; **Supplementary Table 3**). Furthermore, the accumulation of F4/80 immune cells was significantly reduced in both the dorsal root ganglia (**Fig. 2B**) and sciatic nerves (**Fig. 2C**) of *Sele /* mice compared to vincristine-treated C57BL6/J controls at 25 h post first vincristine administration (**Fig. 2B-D**; **Supplementary Table 4**). To determine whether E-selectin deletion confers structural neuroprotection during vincristine exposure, we next quantified intraepidermal nerve fibre density and myelinated fibre integrity by immunohistochemistry and electron microscopy. Notably, no significant differences were detected between saline- and vincristine-treated groups in either genotype. Intraepidermal fibre density in footpads was comparable across C57BL/6J and *Sele /* mice (**Fig. 2D**). Similarly, vincristine did not alter myelinated fibre density (representative images, **Fig. 2F**; quantification, **Fig. 2G**) or the relative frequency distribution of myelinated fibre calibres in caudal nerves (**Fig. 2H**). Together, these findings indicate that vincristine induces sustained mechanical hypersensitivity that is prevented in *Sele /* mice, without causing detectable structural loss of small or myelinated fibres.

**Fig. 2.**
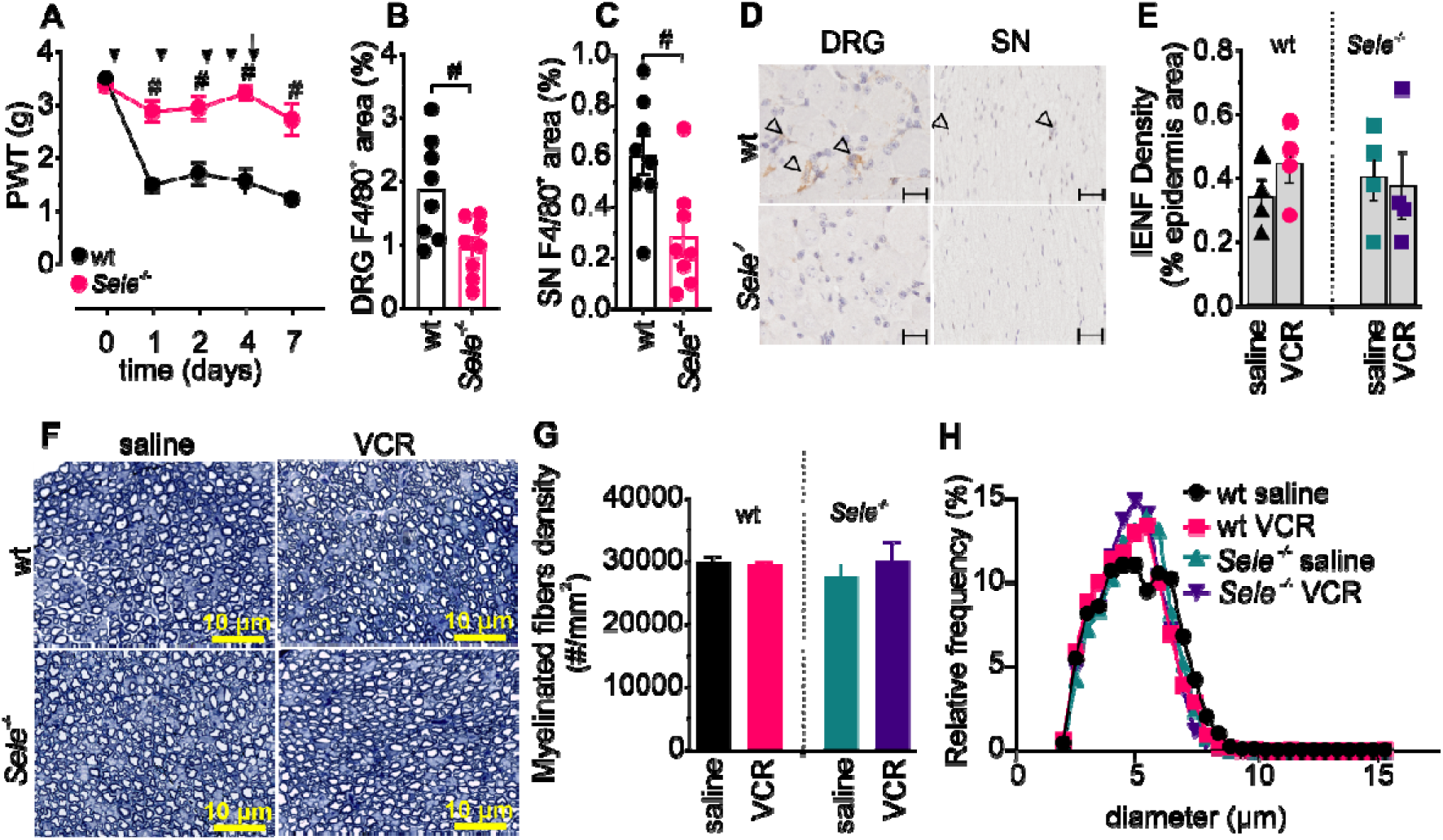
Vincristine induces mechanical hypersensitivity and F4/80 cell accumulation, but not peripheral nerve damage, that is attenuated in Sele□/□mice. **(A)** Sele□/□mice were protected from vincristine-induced mechanical hypersensitivity compared to wt C57BL6/J controls. Mice received vincristine (VCR) for five consecutive days (black arrows), and mechanical hypersensitivity was assessed at baseline (0 h) and on days 1, 2, 4 and 7 post-treatment. **(B,C**) In contrast to wt C57BL6/J mice, Sele□/□mice treated with vincristine were protected from F4/80 cell accumulation in the dorsal root ganglia (DRG) and sciatic nerve (SN) 25 h after the first vincristine administration. **(D)** Representative images of F4/80 immunoreactivity in DRG and SN from wt C57BL6/J and Sele□/□mice. Arrowheads indicate F4/80 areas. Scale bars, 20 µm (DRG) and 10 µm (SN). **(E-H)** Vincristine does not cause structural damage to peripheral nerve fibres. No significant differences in intraepidermal nerve fibre (IENF) density **(E),** myelinated fibre density **(F,G)** or relative myelinated fibre size distribution in caudal nerves **(H)** were observed between saline and vincristine treatments in both wt C57BL6/J and Sele□/□mice. **(F)** Representative images of myelinated fibres in caudal nerves of wt C57BL/6J and Sele□/□mice treated with vincristine or saline. Scale Bar 10 µm. Statistical analyses were performed using repeated-measures two-way ANOVA with Sidak’s multiple-comparison test (A,H) or two-tailed t-tests (B–G). #: P < 0.05; n = 3–8 per group.

### Cell–cell communication roles are reprogrammed by E-selectin perturbation during vincristine treatment

Given the absence of detectable structural nerve damage despite robust mechanical hypersensitivity and immune-cell accumulation (**Fig. 2E–H**), we used spatial transcriptomics of intact DRG sections to identify molecular signatures of neuronal stress and altered multicellular signalling underlying VIPN. This approach also enabled us to determine how genetic deletion or pharmacological blockade of E-selectin alters these responses and to identify the cellular compartments most affected by E-selectin perturbation. DRGs were collected following a single vincristine (0.5 mg kg ¹, i.p.) or saline injection from C57BL/6J mice, *Sele /* mice, and C57BL/6J mice pre-treated with E-selectin-neutralising antibody, and analysed for gene-expression changes and spatial signalling patterns.

First, we performed a pseudobulk analysis of differentially expressed genes (DEGs). Comparing saline-treated C57BL6/J mice to vincristine-treated C57BL6/J mice, we identified 33 upregulated and 44 downregulated genes (**Supplementary Fig. S2**; **Supplementary Table 5A**). Among the most significantly upregulated genes were Map1b (microtubule associated protein 1B) and Hcn2 (hyperpolarisation activated cyclic nucleotide gated potassium and sodium channel 2). Additionally, the most significantly downregulated genes included genes of the haemoglobin family, the Hba-a2 (haemoglobin subunit alpha 2), Hba-a1 (haemoglobin subunit alpha 1) and Hbb-bs (haemoglobin subunit beta).

Enrichment (MsigDB) analysis of DEGs showed robust and conserved stress response in DRG in response to vincristine treatment, characterised by enrichment of mTORC1 signalling, cholesterol and lipid homeostasis, and proteostasis pathways, including the unfolded protein response (UPR), together with activation of p53 and apoptosis modules (**Supplementary Tables 6–7**). At the gene level, this transcriptional backbone was accompanied by induction of injury- and excitability-associated transcripts (**Supplementary Table 5A**), including stress and remodelling markers (e.g. Jun, Map1b) and excitability-linked genes such as Hcn2, alongside downregulation of antigen-presentation genes (e.g. Cd74, H2-Aa, H2-Eb1).

Comparing vincristine-treated C57BL6/J mice to vincristine-treated *Sele*^−/−^ mice, we identified 214 upregulated and 97 downregulated genes (**Supplementary Fig. S2**; **Supplementary Table 5B**), while comparison between vincristine-treated C57BL6/J mice and C57BL6/J mice pre-treated with E-selectin antibody (Esel-ab) prior to vincristine revealed 151 upregulated and 34 downregulated genes (**Supplementary Fig. S2**; **Supplementary Table 5C**). Among the most significantly upregulated genes in both comparisons were *Scand1* (encoding SCAN domain-containing protein 1) and *Hspb1* (encoding heat shock factor binding protein 1), both transcription factors potentially involved in modulating cellular stress responses. Among the downregulated genes in comparisons were *Bst2* (encoding tetherin) and *Ly6a* (encoding stem cell antigen-1), both of which play key roles in the regulation of immune responses. The complete lists of up- and downregulated genes for all conditions are provided in **Supplementary Table 5**. Enrichment analysis of DEGs revealed that perturbation of E-selectin signalling, either by genetic deletion or pharmacological blockade, was associated with prominent downregulation of interferon α/γ response pathways, together with enrichment of hypoxia, epithelial–mesenchymal transition, and immune–vascular signalling programs (**Supplementary Tables 8–11**). Both *Sele^−/−^* and Esel-ab conditions converged on a shared interferon-associated transcriptional module, including *Stat1*, *Stat2*, *Ifitm3*, *Ly6e* and *Bst2*, as well as an extracellular matrix component (*Col1a1*) indicating a conserved E-selectin–sensitive immune state under vincristine challenge. Notably, while the overall transcriptional profiles were overlapping, pharmacological E-selectin blockade exhibited a distinct metabolic signature, with additional enrichment of oxidative phosphorylation, reactive oxygen species, and unfolded protein response pathways (**Supplementary Tables 10–11**), consistent with altered bioenergetic and proteostatic adaptation relative to the knockout condition.

In our transcriptome data, we did not detect increased expression of *Sele* mRNA following vincristine treatment in DRGs. To confirm that vincristine administration is not accompanied with increased E-selectin expression, we also quantified soluble E-selectin (sE-sel) protein in plasma following treatment with vincristine, which failed to detect significant changes in E-selectin expression at the protein level (**Supplementary Fig. S2**). Similarly, *Sele/Hprt* levels analysed using qPCR, remained unchanged in the human endothelial cell line HUVEC following exposure to vincristine (**Supplementary Fig. S2**).

Spotlight analysis identified several neuronal and non-neuronal subtypes within the specific treatment groups (**Supplementary Fig. S2).** Therefore, we next assessed changes in specific cell type-mediated signalling using CellChat analysis ^31^ to further understand how deletion or inhibition of E-selectin may mediate protection from mechanical hypersensitivity in our VIPN model (**Fig. 3, Supplementary Fig. 4, Supplementary Table 12**).

**Fig. 3.**
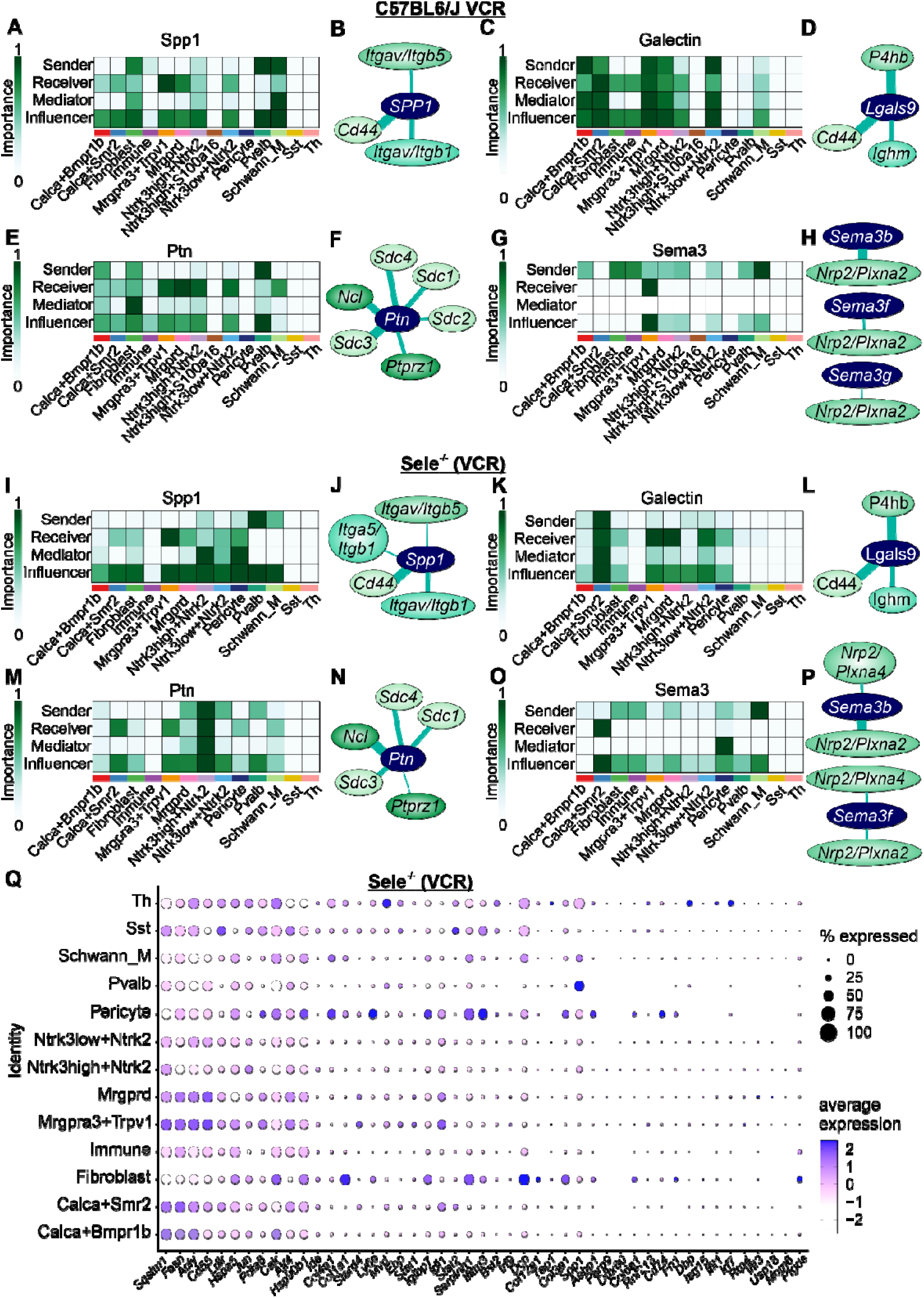
Vincristine reshapes cell-cell communication in dorsal root ganglia through injury-associated signalling pathways. CellChat analysis was used to infer ligand-receptor-mediated communication networks in dorsal root ganglia (DRG) following vincristine treatment, revealing pathway-specific sender, receiver, mediator and influencer roles across neuronal and non-neuronal populations. Heatmaps (**A,C,E,G,I,K,M,O**) show the relative importance of each cell type as senders, receivers, mediators or influencers, with darker green indicating greater contribution. Network schematics (**B,D,F,H,J,L,N,P**) depict dominant ligand–receptor interactions (thickness of the connector line depicts the relative contribution of the L-R pair). (**A-H**) Analysis of Spp1 (**A,B**), galectin (**C,D**), Ptn (**E,F**) and Sema3 (**G,H**) signalling in vincristine-treated C57BL6/J mice. **(I–P)** Corresponding analyses in Sele□/□mice treated with vincristine demonstrate preservation of the same signalling pathways but redistribution of communication roles, with changed immune-cell, stromal and glial populations participation as senders, mediators and influencers across Spp1, galectin, Ptn and Sema3 pathways. **(Q)** Dot plot showing average expression (colour scale) and proportion of expressing cells (dot size) for immune pathway-associated genes across all annotated cell types in Sele□/□(VCR) mice. VCR: vincristine; Sele□/□: E-selectin–deficient mice; DRG: dorsal root ganglion; Schwann_M: myelinating Schwann cells; Calca Bmpr1b and Calca Smr2: peptidergic sensory neuron subtypes expressing Calca with Bmpr1b or Smr2, respectively; Mrgprd: non-peptidergic sensory neurons; Mrgpra3 Trpv1: nociceptive/pruriceptive neurons; Ntrk3^high/+Ntrk2 and Ntrk3^low/+Ntrk2: sensory neuron subsets defined by relative Ntrk3 and Ntrk2 expression; Pvalb: parvalbumin-expressing neurons; Sst: somatostatin-expressing neurons; Th: tyrosine hydroxylase–expressing neurons; Immune: aggregated immune cell populations; Itgav/Itgb: integrin αV/β subunits; Nrp: neuropilin; Plxna: plexin A.

CellChat analysis in C57BL6/J mice treated with vincristine revealed a communication network in which immune cells, fibroblasts, pericytes, and Schwann cells act as dominant senders across multiple injury-associated pathways following vincristine treatment (C57BL6/J (VCR); **Supplementary Table 12)** including Spp1 (osteopontin), Galectin (β-galactoside–binding lectins), Sema3 (semaphorin 3), Kit (stem cell factor receptor), Nrg (neuregulin), Nt (neurotrophins), Nts (neurotensin) and Fgf (fibroblast growth factors) that collectively regulate inflammatory adhesion, immune modulation, axon guidance, growth factor activation, glial support, neuronal survival, pain signalling and tissue remodelling, while sensory neuron subtypes function primarily as receivers (**Fig. 3A,C,E,G; Supplementary Fig. S3**). In particular, immune and support cell populations showed high sender and influencer scores for Spp1, Galectin and Sema3, indicating that these cells not only initiate signalling but also strongly shape network structure.

For Spp1 signalling, immune, fibroblast, Schwann cells and Pvalb neurons were the principal senders and influencers, expressing Spp1 (ligand), while nociceptive and peptidergic neurons (Calca Bmpr1b, Calca Smr2, Mrgpra3 Trpv1, Ntrk3^high/+Ntrk2) acted as dominant receivers, expressing CD44 and integrin receptors (Itgav/Itgb1, Itgav/Itgb5) (**Fig. 3A, B, Supplementary Table 12**). Fibroblasts and Schwann cells additionally showed elevated mediator scores, consistent with their role in relaying immune-derived signals through the neurovascular niche.

Galectin signalling was driven by neuronal senders (Calca Bmpr1b, Calca Smr2, Mrgpra3 Trpv1, Mgprd) expressing Lgals9, with immune, Schwann cell, fibroblast and some neuronal populations as primary receivers via CD44 and Ighm receptors (**Fig. 3C, D, Supplementary Table 12**). Ptn signalling was dominated by Calca Bmpr1b neurons, Pvalb neurons and fibroblast as major sender via Ptn sending signals towards a mixed population (**Fig. 3E, F, Supplementary Table 12**). Sema3 signalling showed strong non-neuronal sender and influencer activity, with Sema3b/f/g ligands produced by immune cells, fibroblasts and Schwann cells, while Mrgpra3 Trpv1 neurons served as the dominant receivers via Nrp2/Plxna2 and Nrp2/Plxna4 complexes (**Fig. 3G,H, Supplementary Table 12**). Collectively, this data suggests that vincristine establishes a network in which immune and stromal cells act as upstream signal initiators and network shapers, delivering injury-associated cues to neuronal targets.

In E-selectin-deficient animals treated with vincristine (Sele^−/−^ (VCR)), CellChat analysis demonstrated a marked redistribution of communication roles rather than pathway loss (**Fig. 3I–P; Supplementary Table 12, Supplementary Fig. S3**). For Spp1, compared to C57BL6/J (VCR), ligand expression in Sele^−/−^ (VCR) was reduced in immune cells, fibroblasts and Schwann cell acting as senders, while neurons somewhat retained receiver capacity through CD44 and integrins, but with increased influencer coupling (**Fig. 3I,J, Supplementary Table 12**). Galectin signalling showed a reduction in almost all cell types, with Calca Smr2 dominating the signalling pathway (**Fig. 3K,L, Supplementary Table 12**). Ptn signalling in Sele^−/−^ (VCR) showed decreased sender and mediator roles for immune cells and fibroblasts, while Schwann cell and pericytes signals increased, dominated by Ntrk3^high/+Ntrk2 neurons. Ptn signalling pathway was modulated via Ptprz1 ligand and syndecan receptors (Sdc1–4) (**Fig. 3M,N, Supplementary Table 12**). In Sema3, non-neuronal sender and influencer scores were reduced, and signalling was redistributed among non-neuronal populations (**Fig. 3O,P, Supplementary Table 12**).

Pharmacological blockade of E-selectin (C57BL/6J (Esel-ab + VCR)) produced a communication architecture that overlapped with, but extended beyond, the knockout condition (**Supplementary Fig. S3, Supplementary Table 12**). In this setting, immune cells, fibroblasts and Schwann cells showed further reduction in sender and influencer roles across injury-associated pathways.

Additionally, we observed a striking suppression of genes involved in inflammatory pathways in *Sele^−/−^* treated with vincristine (**Supplementary Tables 8-9**) suggesting that E-selectin may act as an upstream driver of neuroinflammation in VIPN, potentially by modulating inflammatory processes, such as interferon-related immune activation. To further investigate this, we performed gene expression plot of immune signalling associated genes in *Sele^−/−^* to determine cell populations that are primarily responsible for the shift in inflammation-associated genes (**Fig. 3Q**). Interferon-responsive transcripts, including *Stat1*, *Stat2, Ifitm3, Bst2, Ly6e* and *Isg15*, were supressed predominantly within immune cell clusters (**Fig. 3Q**). Fibroblasts and pericytes displayed strong expression of extracellular matrix and remodelling genes (*Col1a1, Col3a1, Col4a1, Col12a1, Dcn, Fn1*), while Schwann cells co-expressed immune-responsive (*Ifitm3, Ly6e*) and matrix-associated (*Col4a1, Serpinh1*) transcripts, positioning them at the interface of immune–stromal signalling. In contrast, sensory neuron subtypes (*Calca Bmpr1b*, *Calca Smr2*, *Mrgpra3 Trpv1*, *Mrgprd*, *Ntrk3*-defined populations) exhibited minimal interferon or extracellular matrix gene expression but retained stress-associated transcripts (*Jun, Atf4, Hspa5*).

In summary, under vincristine treatment, dorsal root ganglia exhibit a transcriptional and cell-cell communication profile marked by activation of stress, interferon, and immune–stromal signalling programs, with immune cells, fibroblasts, pericytes, and Schwann cells acting as dominant senders and influencers across multiple injury-associated pathways while sensory neurons function primarily as receivers. In contrast, genetic deletion or pharmacological blockade of E-selectin preserves these pathways but redistributes communication roles, reducing immune-cell dominance, altering interferon- and metabolism-associated gene expression, and increasing the involvement of stromal and glial populations in signal mediation.

### E-selectin-induced mechanical hypersensitivity is mediated by immune cells

Spatial transcriptomic analyses indicated that E-selectin may influence vincristine-induced mechanical hypersensitivity through modulation of immune cell activity and inflammatory signalling, extending beyond its canonical role as an endothelial adhesion molecule. To directly assess this possibility, we administered recombinant E-selectin locally into the hind paw of C57BL6/J mice (intraplantar, i.pl.) and quantified mechanical paw withdrawal thresholds. This approach permitted investigation of the direct effects of E-selectin on peripheral immune or neuronal cells, bypassing the vascular endothelium and excluded confounding effects from endothelial adhesion.

Remarkably, i.pl. injection of E-selectin induced robust mechanical hypersensitivity, detectable as early as 1 hour post-injection and persisting at 4, 24, and 48 hours (**Fig. 4A; Supplementary Table 13**). In contrast, saline or IgG2b controls failed to induce hypersensitivity (**Fig. 4A; Supplementary Table 13**). Notably, E-selectin injection did not produce heat hyperalgesia (**Fig. 4B**). This mechanical hypersensitivity was accompanied by a significant increase in F4/80 cell area in injected footpads compared to saline or IgG2b (**Fig. 4C**), with representative immunohistochemistry shown in **Figure 4D**.

**Figure 4.**
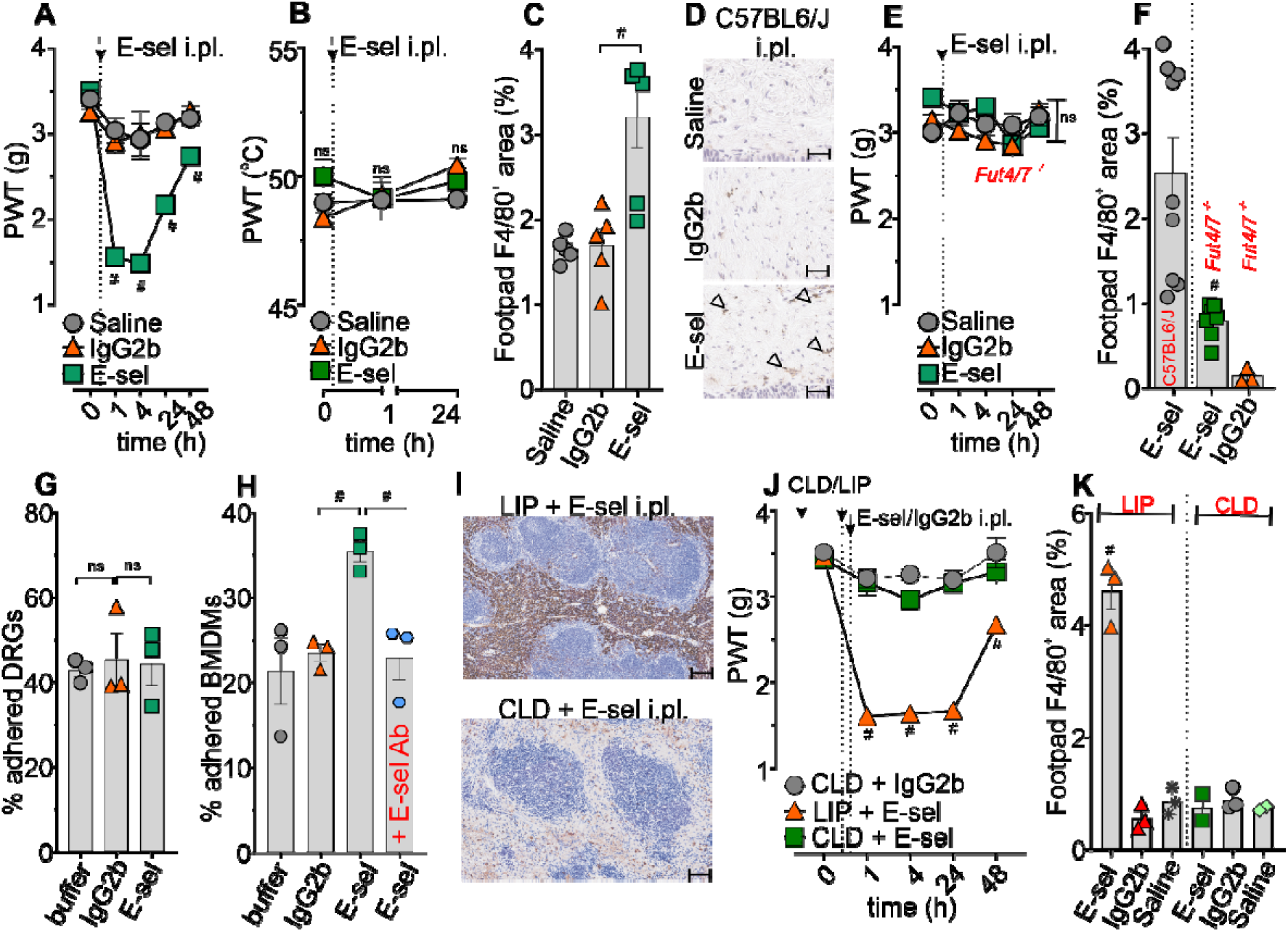
E-selectin-induced mechanical hypersensitivity is mediated by immune cells. (**A**) Local intraplantar (i.pl.) injection of E-selectin – but not saline or IgG2b – into the hind paw of C57BL6/J mice induces significant mechanical hypersensitivity, (**B**) but not thermal hypersensitivity, and is (**C**) accompanied by an increase in F4/80 cells in the injected footpads. **(D)** Representative images of F4/80 cells in the E-selectin, IgG2b and saline-injected food pads. Scale bar = 10 µm. (**E**) Fut4/7□/□mice are protected from E-selectin-induced mechanical hypersensitivity and (**F**) the E-selectin-induced increase in F4/80 cell area. (**G–H**) Adhesion of dissociated DRG neurons (G) and BMDMs (H) to surfaces coated with recombinant E-selectin, IgG control or TBS (5 μg/mL). Cells were allowed to adhere for 1 h before non-adherent cells were removed by washing. (**G**) E-selectin does not increase DRG neuron adhesion, but (**H**) significantly increases BMDM adhesion, which is significantly reduced by pre-treatment with an E-selectin–blocking antibody (E-sel Ab). **(I)** Representative immunohistochemical images of F4/80 cells in the spleens of C57BL6/J mice pretreated with clodronate (CLD) or liposome control (LIP) and injected i.pl. with E-sel, demonstrating effective depletion of F4/80 cells. Scale bar = 10 µm. (**J**) Pretreatment with liposomal clodronate (CLD) protects C57BL6/J mice from E-selectin-induced mechanical hypersensitivity. (**K**) CLD pretreatment also significantly reduces the increase in F4/80 cell area in the footpad following i.pl. E-selectin injection. Statistical significance (#: P < 0.05) was determined using one-way ANOVA for C, F, G, H, K (n=3-9) or repeated measures two-way ANOVA for A, B, E, J (n = 6-9).

To confirm ligand specificity, i.pl. E-selectin injection was repeated in Fut4/7□/□mice, which lack functional α(1,3)-fucosyltransferases required for the synthesis of E-selectin ligands^36^. Fut4/7^□/□^ mice did not develop mechanical hypersensitivity following E-selectin injection (**Fig. 4E**), and their footpad F4/80 area was significantly reduced compared to C57BL6/J-E-selectin treated animals and indistinguishable from IgG2b controls (**Fig. 4F**).

To define the cellular mediators of this hypersensitivity, adhesion assays were performed using dorsal root ganglia (DRG) neurons and bone marrow–derived macrophages (BMDMs) seeded onto E-selectin–coated surfaces. DRG adhesion was unaffected by E-selectin (**Fig. 4G**), suggesting little to no direct interaction between neurons and E-selectin. In contrast, BMDMs exhibited significantly increased adhesion to E-selectin compared to controls, which was abolished by E-selectin–blocking antibodies (**Fig. 4H**). These findings suggest that E-selectin primarily interacts with immune cells, such as macrophages, rather than neurons.

To further confirm the specific role of immune-mediated mechanisms in E-selectin–induced mechanical hypersensitivity, phagocytic cells were depleted with liposomal clodronate (CLD), with depletion confirmed in spleens (**Fig. 4I**). C57BL6/J mice pretreated with CLD or control liposomes (LIP) then received i.pl. injections of E-selectin, IgG2b, or saline (**Fig. 4J**). While LIP-pretreated mice developed sustained mechanical hypersensitivity following E-selectin injection, CLD-pretreated mice failed to develop hypersensitivity (**Fig. 4J, Supplementary Table 14**). No hypersensitivity was observed in saline or IgG2b controls (**Supplementary Fig. S4A**). Consistent with this, CLD pretreatment significantly reduced F4/80 cell area in injected footpads (**Fig. 4K**), with representative images shown in **Supplementary Fig. S4B**.

### E-selectin adhesion enhances interleukin-1β release in vincristine-treated BMDMs

We next investigated whether E-selectin adhesion alters vincristine-induced cytokine and chemokine release in BMDMs to determine whether the role of E-selectin in development of vincristine-induced neuropathy is restricted to its known canonical cell adhesion function.

As expected and previously reported^8^, LPS + VCR treatment significantly increased (∼40-fold) the secretion of interleukin-1β (IL-1β) in unadhered BMDMs (**Fig. 5A**). Interestingly, we also detected a significant increase of interleukin-18 (IL-18, **Fig. 5B**), interleukin-10 (IL-10, **Fig. 5C**), granulocyte colony-stimulating factor (G-CSF, **Fig. 5D**), interleukin-12 subunit p40 (IL-12p40, **Fig. 5E**), interleukin-6 (IL-6, **Fig. 5F**) and the chemokine C-X-C motif chemokine ligand 1 (CXCL1, **Fig. 5G**) (values of the cytokines in test and control conditions are shown in **Supplementary Table 15**). No significant changes were observed for tumour necrosis factor alpha (TNF-α), transforming growth factor beta 1 (TGF-β1), interleukin-23 (IL-23), interleukin-12 p70 heterodimer (IL-12p70), C-C motif chemokine ligand 17 (CCL17; also known as thymus and activation-regulated chemokine, TARC), or C-C motif chemokine ligand 22 (CCL22; also known as macrophage-derived chemokine, MDC) (**Supplementary Table 15**).

**Fig. 5.**
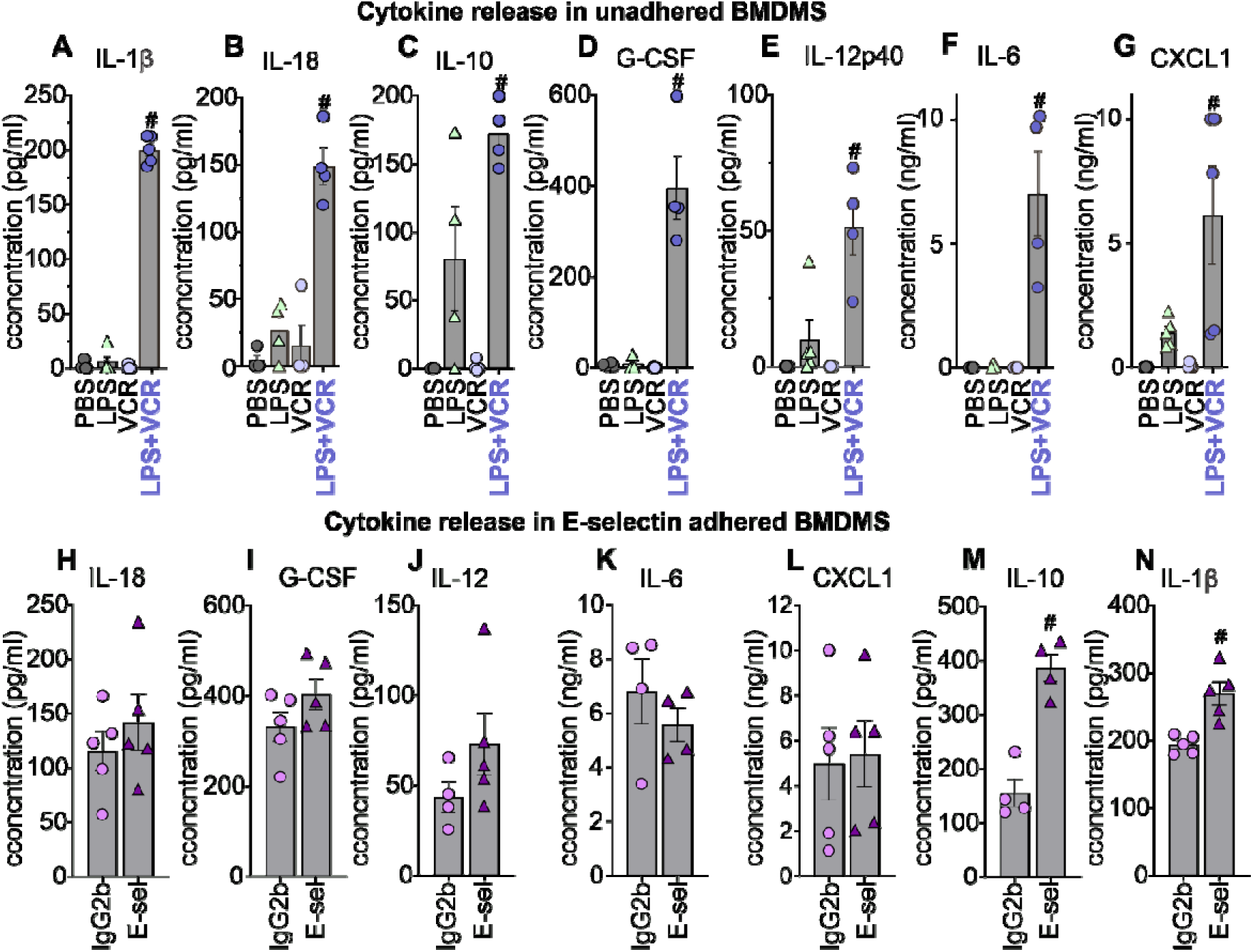
E-selectin adhesion specifically enhances interleukin-1β release in vincristine-treated BMDMs. (**A-G**) BMDMs were primed with LPS (10 ng/mL, 3 h) followed by vincristine (VCR; 100 μM, 6 h). Release of IL-1β (**A**), IL-18 (**B**), IL-10 (**C**), G-CSF (**D**), IL-12p40 (**E**), IL-6 (**F**) and CXCL1 (**G**) was increased in LPS+VCR-treated BMDMs compared with PBS, LPS or VCR controls. (**H–N**) BMDMs were adhered to IgG2b- or E-selectin-coated surfaces (5 μg/mL) and treated with LPS (10 ng/mL, 3 h) followed by VCR (100 μM, 6 h). Release of IL-18 (**H**), G-CSF (**I**), IL-12 (**J**), IL-6 (**K**) and CXCL1 (**L**) did not differ between IgG2b- and E-selectin-adhered BMDMs, whereas IL-10 (**M**) and IL-1β (**N**) release was increased ∼2.5- and ∼1.4-fold, respectively, following E-selectin adhesion. Statistical significance was determined using one-way ANOVA with Dunnett’s multiple comparisons test, comparing LPS+VCR with LPS alone (A–G) or E-selectin- with IgG2b-adhered BMDMs (H–N) (n = 4 per group; #P < 0.05).

The adhesion of BMDMs to E-selectin did not significantly enhance the release of interleukin-18 (IL-18, **Fig. 5H**), granulocyte colony-stimulating factor (G-CSF, **Fig. 5I**), interleukin-12 subunit p40 (IL-12p40, **Fig. 5J**), interleukin-6 (IL-6, **Fig. 5K**), and C-X-C motif chemokine ligand 1 (CXCL1, **Fig. 5L**, (**Supplementary Table 16**). However, BMDMs adhered to E-selectin and treated with LPS + VCR released approximately 2.5-fold more IL-10 (**Fig. 5M**) and 1.4-fold more IL-1β (**Fig. 5N**) compared to IgG2b controls. Notably, only IL-1β release, but not release of other cytokines, was significantly inhibited by pre-treatment with the NLRP3 inflammasome inhibitor MCC950 (10 µM)^13^, indicating a specific inflammasome-dependent mechanism (values of the cytokines in test and control conditions, including LPS, nigericin and inhibition with MCC950, are shown in **Supplementary Table 16**).

### E-selectin adhesion enhances NLRP3 signalling in vincristine-induced neuropathy

Given the established role of the NLRP3 inflammasome in vincristine-induced IL-1β release, we next investigated whether the effects of E-selectin on IL-1β release observed above in our study are specifically mediated through enhanced NLRP3 inflammasome activation. To assess this, we employed ASC-citrine reporter BMDMs, which allow real-time quantification of ASC speck formation (**Figure 6A**) - a direct readout of NLRP3 inflammasome activation.

**Figure 6.**
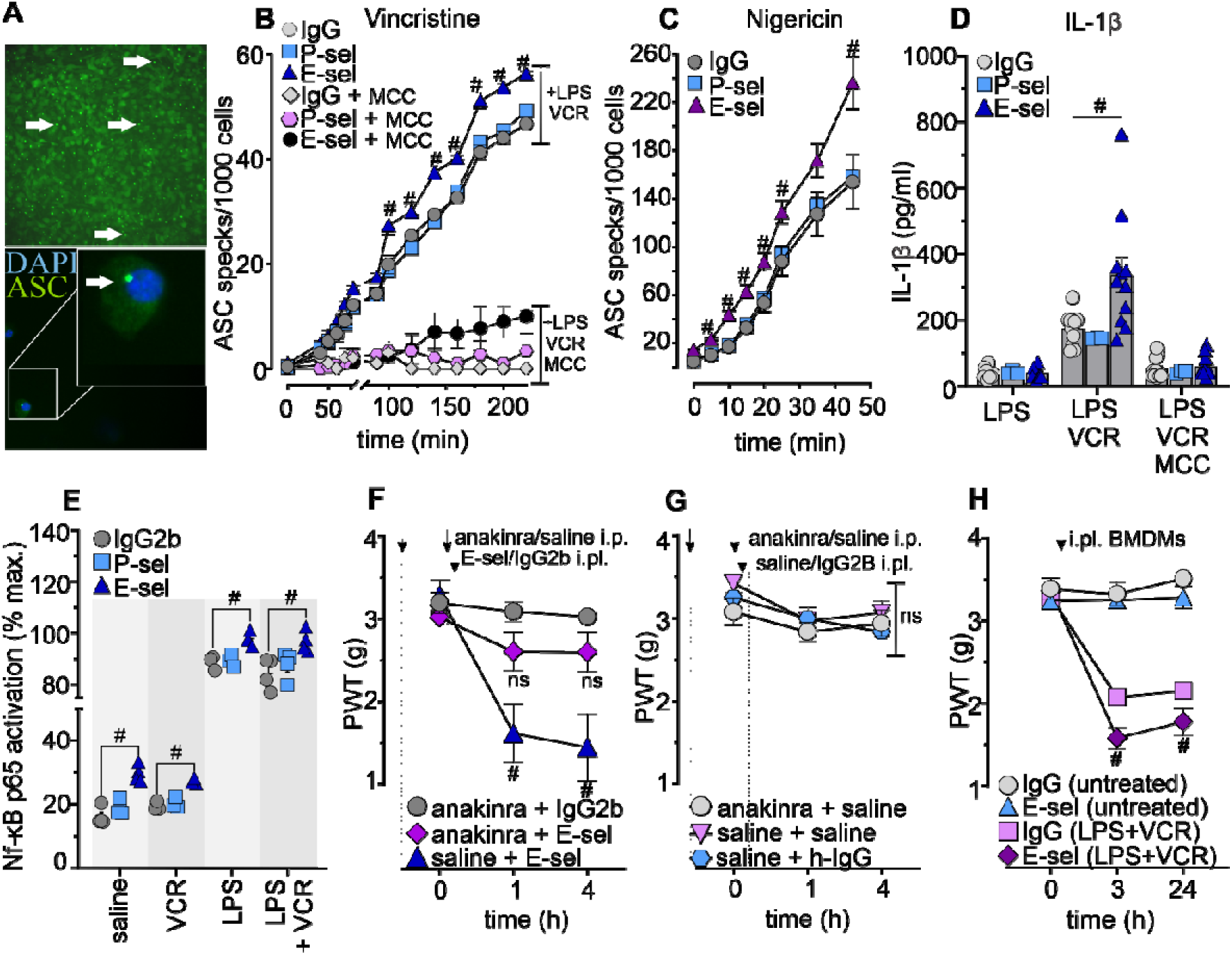
E-selectin functions as a non-canonical signalling amplifier of NF-κB– dependent NLRP3 activation and IL-1β release, driving vincristine-induced mechanical hypersensitivity. BMDMs were adhered to E-selectin-, P-selectin- or IgG-coated surfaces (5 μg/mL), primed with LPS (10 ng/mL, 3 h), and subsequently treated with VCR (100 μM, 6 h) or nigericin (5 μM, 45 min), as indicated. Where indicated, cells were pre-treated with MCC950 (10 μM, 30 min) before stimulation. **(A)** Representative images of ASC specks (green; white arrows) in bone marrow-derived macrophages (BMDMs) treated with lipopolysaccharide (LPS, 10 ng/mL) and vincristine (VCR, 100 µM). Nuclei are stained with DAPI (blue). Top: 10× magnification; bottom:60× magnification. **(B,C)** Quantification of ASC specks over time in ASC–citrine BMDMs adhered to E-selectin (E-sel), IgG2b, or P-selectin (P-sel) following treatment with LPS + VCR (**B**) or LPS + nigericin (**C**). E-selectin adhesion significantly increased ASC speck formation compared to controls. This effect was fully blocked by pre-treatment with the NLRP3 inhibitor MCC950 (10 µM) and was mirrored by IL-1β release (**D**). (**E**) Adhesion to E-selectin significantly increased the proportion of active p65 NF-κB across all treatment conditions (saline, vincristine, LPS, and LPS + vincristine) compared with IgG2b controls, whereas P-selectin did not induce comparable NF-κB activation. (**F**) Intraplantar (i.pl.) injection of E-selectin causes IL-1β-mediated mechanical hypersensitivity that is prevented by the IL-1 receptor antagonist anakinra (100 mg/kg; ip.). (**G**) Intraplantar administration of anakinra or saline prior to PBS or IgG2b i.pl. injection does not cause mechanical hypersensitivity. **(H)** Intraplantar injection of BMDMs treated with LPS + VCR and adhered to E-selectin induced significantly greater mechanical hypersensitivity compared to IgG2b-adhered controls. Injection of untreated BMDMs adhered to either E-sel or IgG2b had no effect. Statistical significance (#: P < 0.05) was determined using repeated measures two-way ANOVA (n = 6). Statistical significance (n=6) was determined by repeated-measures two-way (**B,C,F,G,H**) or one-way ANOVA (**D,E**) with Dunnett’s multiple comparisons test (#P < 0.05).

BMDMs adhered to E-selectin and treated with lipopolysaccharide (LPS, 10 ng/mL) and vincristine (VCR, 100 µM) formed ASC specks more rapidly and in significantly greater numbers compared to cells adhered to IgG or P-selectin (**Fig. 6B; Supplementary Table 17**). A similar enhancement in ASC speck formation was observed when E-selectin– adhered BMDMs were treated with LPS (10 ng/mL) and nigericin (5 µM) (**Fig. 6C; Supplementary Table 18**). In both cases, ASC speck formation was completely abrogated by pre-treatment with MCC950 (10 µM), a selective NLRP3 inflammasome inhibitor, confirming that the observed effects were NLRP3-dependent (**Fig. 6B; Supplementary Fig. S5A-B; Supplementary Table 17-18**).

This increase in ASC speck formation was accompanied by significantly elevated secretion of IL-1β from E-selectin-adhered BMDMs in response to both LPS + VCR and LPS + nigericin (**Fig. 6D; Supplementary Fig. S5C; Supplementary Table 19**). By contrast, treatment with LPS, MCC950, nigericin, vincristine, or PBS alone, in the context of E-selectin adhesion, did not induce ASC speck formation (**Supplementary Fig. S5; Supplementary Table 17-18)** or IL-1β release **(Supplementary Table 19**).

To assess whether this response was specific to vincristine, BMDMs adhered to E-selectin, IgG, or buffer, were also treated with LPS in combination with other chemotherapeutic agents, including doxorubicin (DOX, 50 µM) and cyclophosphamide (CYC, 100 µM). Although low levels of IL-1β release were detected under these conditions, E-selectin adhesion had no significant effect, indicating that the potentiating effect of E-selectin is specific to vincristine (**Supplementary Figure S5D-E, Supplementary Table 20**).

Collectively, these data suggest that E-selectin does not simply function as a cellular adhesion molecule, but has additional, non-canonical functions as an amplifier of NLRP3 signalling and IL-1β release. Given that NF-κB signalling provides a critical priming step for NLRP3 activation, and that E-selectin engagement enhances NF-κB activity in leukocytes^37^, we hypothesised that E-selectin and vincristine pathways functionally converge on NF-κB to potentiate inflammasome activation and IL-1β release. Indeed, E-selectin adhesion significantly increased the proportion of active p65 NF-κB across all treatment conditions, compared to IgG2b controls (**Fig. 6E**).

Next, to determine whether these non-canonical signalling effects also contribute to development of vincristine-induced neuropathy *in vivo*, we assessed the effect of the IL-1 receptor antagonist anakinra on mechanical paw withdrawal thresholds following i.pl. injection of E-selectin (**Fig. 6F-G**). Interestingly, anakinra significantly attenuated E-selectin–induced mechanical hypersensitivity compared to saline (**Fig. 6F, Supplementary Table 21**), while no hypersensitivity was observed in control groups (**Fig. 6G, Supplementary Table 21**).

### E-selectin amplifies NLRP3-dependent IL-1β release from vincristine-stimulated macrophages in vitro, while IL-1 receptor signalling contributes to E-selectin-induced mechanical hypersensitivity in vivo

We previously demonstrated that intraplanar injection of BMDMs treated with LPS and vincristine into the hind paw of mice induces local mechanical allodynia resembling VIPN^8^. To assess whether the non-canonical signalling effects of E-selectin – leading to enhanced NLRP3 activation and IL-1β release in response to vincristine – also contribute to VIPN symptoms *in vivo*, we injected LPS + VCR–treated BMDMs adhered to E-selectin into the hind paw of mice. As expected, E-selectin–adhered BMDMs treated with LPS + VCR induced significantly more severe mechanical hypersensitivity than IgG2b-adhered cells (**Fig. 6H**). In contrast, intraplantar injection of untreated BMDMs adhered to either E-selectin or IgG2b had no effect on mechanical sensitivity.

Overall, our results position E-selectin as a putative anti-allodynic target for treatment of VIPN. We show that in addition to enhanced immune cell infiltration into neuronal tissue, immune cell adhesion to E-selectin directly enhances NLRP3-mediated IL-1β release via enhanced Nf-kB activation. We thus show a non-canonical signalling role of E-selectin in the pathogenesis of VIPN for the first time.

## Discussion

Vincristine-induced peripheral neuropathy (VIPN) has traditionally been understood as a consequence of direct axonal toxicity and microtubule disruption. Our findings instead suggest a broader view: VIPN behaves as a neuroimmune network disorder in which interactions between vascular adhesion molecules and immune cells reorganise communication within the dorsal root ganglia (DRG) before any clear structural nerve degeneration becomes apparent. Within this framework, E-selectin should not be considered simply as an adhesion molecule involved only in immune cell recruitment. Rather, it functions as a higher-level regulator that coordinates immune cell trafficking and amplifies inflammatory signalling within peripheral nerves.

By examining representative members of major vascular adhesion-molecule families, we identified a selective pathogenic signature in VIPN. Of the leukocyte adhesion molecules tested, only E-selectin and VCAM-1 contributed to vincristine-induced mechanical hypersensitivity and accumulation of F4/80 immune cells, while blockade of ICAM-1, PECAM-1 or P-selectin did not provide protection (**Fig. 1**). This indicates that the neuropathy is not driven by broad endothelial activation, but instead depends on a specific adhesion axis. Importantly, the translational potential of targeting this axis differs between molecules. VCAM-1-directed therapies have encountered safety and implementation barriers as endothelial-surface VCAM-1 is important for leukocyte tissue homing. On the other hand, E-selectin is largely redundant for this purpose and E-selectin inhibitors such as Uproleselan are well-tolerated in oncology trials ^38–40^. This difference makes E-selectin a particularly actionable target within the vascular–immune interface.

These findings extend our previous demonstration that macrophage depletion, genetic or pharmacological inhibition of NLRP3, and IL-1 receptor antagonism protect against vincristine-induced neuropathy^8,14^. Here, we sought to define how E-selectin is mechanistically linked to this established macrophage–NLRP3–IL1β pathway and to sensory hypersensitivity. A critical component of this question was whether E-selectin acts directly on sensory neurons or indirectly through immune cells. Intraplantar administration exposed peripheral sensory terminals directly to E-selectin while permitting assessment of a local, unilateral mechanical response. However, E-selectin promoted macrophage but not DRG neuron adhesion, and E-selectin-induced hypersensitivity was attenuated by phagocyte depletion and IL-1 receptor antagonism. These findings support a predominantly immune-mediated mechanism rather than direct sensitisation of sensory neurons.

Together with the protection afforded by E-selectin blockade or deletion in the systemic vincristine model, our findings position E-selectin at the point of convergence between immune-cell recruitment and inflammatory signalling in VIPN. E-selectin facilitates macrophage accumulation within peripheral neuronal tissues while also enhancing vincristine-induced NF-κB activation, NLRP3 inflammasome assembly and IL1β release. E-selectin thereby couples immune-cell recruitment and activation to IL1R-dependent sensory hypersensitivity.

Notably, the administration of vincristine induced persistent mechanical hypersensitivity without loss of intraepidermal or myelinated nerve fibres (**Fig. 2**). This finding reframes VIPN in our model as a condition induced by functional neuroimmune reprogramming rather than primary structural degeneration. Spatial transcriptomic analyses of DRGs revealed a consistent stress- and immune-associated transcriptional programme characterised by activation of mTORC1, engagement of unfolded protein response pathways, and induction of excitability-related genes including *Jun* and *Hcn2* (**Fig. 3**). These transcriptional changes did not occur in isolation. Instead, they were embedded within a reorganised intercellular communication landscape in which immune cells, fibroblasts, pericytes and Schwann cells acted as dominant signal senders across SPP1, Galectin, PTN, SEMA3 and growth factor pathways, while sensory neurons predominantly functioned as receivers. In this configuration, vincristine establishes a hierarchical inflammatory structure in which neuronal dysfunction is shaped by signalling from surrounding non-neuronal cells.

Importantly, perturbation (inhibition or genetic deletion) of E-selectin did not eliminate this communication network but instead altered its structure (**Fig. 3**). The dominance of immune cells as signalling hubs was reduced, and stromal (non-neuronal) populations assumed greater mediator roles. These findings indicate that E-selectin does not simply act as a leukocyte homing molecule induced following vincristine-mediated damage; rather, E-selectin-mediated signalling amplifies and stabilises inflammatory network states. In this sense, E-selectin may function as a regulatory gate that determines whether chemotherapeutic stress resolves adaptively or progresses into sustained nociceptive signalling.

Although the genetic studies used age-matched C57BL/6J rather than littermate controls, the protective phenotype observed in *Sele*^−/−^ mice was independently recapitulated by pharmacological E-selectin blockade in C57BL/6J mice, supporting an E-selectin-dependent mechanism while reducing the likelihood that the findings are attributable solely to genetic-background effects.

Mechanistically, E-selectin exerts two converging effects. First, in its canonical role, it promotes F4/80 immune cell recruitment into neuronal tissues providing the cellular foundation for inflammation (**Fig. 1**). Second, in a non-canonical role, E-selectin-mediated adhesion enhances NF-κB activation and NLRP3 inflammasome–dependent IL-1β release in macrophages, selectively amplifying cytokine production during vincristine exposure (**Fig. 5-6**). Together, these findings define a feed-forward inflammatory loop that operates upstream of neuronal sensitisation. Furthermore, intraplantar administration of E-selectin was sufficient to induce phagocyte-dependent mechanical hypersensitivity *in vivo* via IL1R signalling, confirming that its pro-nociceptive effects arise primarily through immune-tissue environment interactions rather than direct neuronal activation (**Fig. 4**).

Additionally, we administered recombinant E-selectin locally into the paw (**Fig. 4**) to test whether non-canonical E-selectin signalling is sufficient to drive neuroimmune responses that drive mechanical hypersensitivity, independent of its canonical endothelial adhesion role. In this context, E-selectin likely engages its ligands on resident immune cells to promote inflammatory signalling.

Primary sensory neurons innervating the hind paw have their cell bodies in the DRG, axons within peripheral nerves including the sciatic nerve, and sensory terminals in the plantar skin. Analysis of these tissues therefore examines distinct compartments along the same peripheral sensory pathway. Systemic vincristine increased F4/80-positive immune-cell accumulation in both the DRG and sciatic nerve, demonstrating inflammation surrounding the somatic and axonal compartments of peripheral sensory neurons. Although we did not quantify immune-cell accumulation in the footpad following systemic vincristine, repeated systemic treatment caused no detectable loss of intraepidermal nerve fibres. Moreover, previous work directly comparing systemic and intraplantar vincristine demonstrated that both routes produce mechanical allodynia through a shared NLRP3–IL1β-dependent mechanism, with hypersensitivity attenuated by Nlrp3 deletion or pharmacological NLRP3 inhibition with MCC950 ^8,18^.

The intraplantar E-selectin paradigm used here was designed to determine whether E-selectin is locally sufficient to induce immune-dependent hypersensitivity within the peripheral territory innervated by DRG neurons. Intraplantar administration restricts E-selectin exposure to one hind paw, allowing its local effects to be distinguished from its systemic endothelial functions. Local E-selectin administration caused unilateral mechanical hypersensitivity and increased F4/80-positive immune-cell accumulation in the injected footpad, with both responses dependent on functional E-selectin ligands and phagocytic cells. Together with the preferential adhesion of bone marrow-derived macrophages rather than DRG neurons to E-selectin, these findings demonstrate the local sufficiency of E-selectin-dependent immune signalling. This paradigm should therefore be interpreted as a mechanistic sufficiency experiment rather than as an anatomical reproduction of systemic VIPN

Collectively, these results support a model in which VIPN reflects dysregulated neuroimmune network dynamics rather than a simple linear axonopathy. Within this network, E-selectin occupies a central position, linking vascular activation and immune cell recruitment to inflammasome activation which in turn determine both the strength and persistence of pain signalling. Because E-selectin acts upstream of IL-1β and NLRP3, it represents a higher-order control point capable of influencing multiple downstream inflammatory pathways simultaneously.

This conceptual shift has clinical implications for patients. The risk of VIPN-induced morbidity remains a major reason for vincristine dose reduction or chemotherapy discontinuation, yet current treatment options are largely symptomatic. Duloxetine provides only modest benefit, and no established preventive therapies exist^41^. Targeting E-selectin offers a mechanistically grounded approach aimed at modifying disease progression rather than merely masking symptoms. The prior clinical development of E-selectin inhibitors enhances translational feasibility and raises the possibility of protecting neuronal function without compromising anti-tumour efficacy. Uproleselan (GMI-1271, small molecule E-selectin inhibitor) has been evaluated in multiple AML trials, including safety phase I/II studies (NCT02306291, NCT03381795), phase III trial in relapsed/refractory AML (NCT03616470), salvage therapy for relapsed/refractory AML (NCT03616470) and for newly diagnosed older AML patients (NCT03701308, NCT04848974) showing generally favourable tumour adverse event profiles. Nevertheless, careful evaluation in human cohorts is required to determine whether targeting this vascular–immune axis can provide effective analgesia without impairing immune competence or vincristine chemotherapy effectiveness.

Several limitations should be considered. The study relies predominantly on murine models, which may not fully replicate the complexity and heterogeneity of VIPN in patients. Spatial transcriptomic analyses were performed on a limited number of biological replicates. Behavioural assessments focused mainly on mechanical hypersensitivity, and the diversity of tissue-resident macrophage populations was not directly characterised. In addition, the long-term systemic consequences of sustained E-selectin inhibition in tumour-bearing hosts remain elusive. Future studies should examine how this adhesion-dependent network evolves over time, whether similar mechanisms operate across different chemotherapeutic agents, and how cell-type–specific manipulation of E-selectin signalling reshapes nociceptive circuitry.

Moreover, the present study did not directly compare the cellular composition of inflammatory infiltrates across DRG, peripheral nerve and footpad compartments. Accordingly, the intraplantar E-selectin model should be interpreted as a mechanistic sufficiency model rather than as an anatomical reproduction of systemic VIPN. Although previous genetic and pharmacological studies establish NLRP3 as a critical mediator of VIPN ^8,14^, NLRP3 activation was not directly measured in vivo following local E-selectin administration in the present study. Thus, our in vivo data establish dependence on IL-1 receptor signalling, whereas the specific amplification of NLRP3 activation by E-selectin is demonstrated directly in macrophages.

Additionally, ICAM-2 was not examined in the present study. Unlike inducible ICAM-1 and E-selectin, ICAM-2 is constitutively expressed on endothelial cells and has context-dependent roles in leukocyte transmigration; inflammatory cytokines can reduce endothelial ICAM-2 expression ^42,43^. Thus, although our results identify E-selectin and VCAM-1 among the adhesion molecules tested, they do not exclude contributions from ICAM-2 or other endothelial adhesion pathways.

## Conclusion

In conclusion, our findings show that E-selectin plays a central role in coordinating the immune and inflammatory processes that drive vincristine-induced peripheral neuropathy. Rather than simply treating symptoms of VIPN, targeting E-selectin offers the possibility of intervening earlier in the disease process by disrupting the inflammatory pathways that sustain chemotherapy-induced neuropathy.

**Table 1.** Antibodies used for in vivo blockade and tissue staining.

| Application | Antibody or control | Clone | Host/isotype | Supplier | Catalogue no. | RRID |
| --- | --- | --- | --- | --- | --- | --- |
| E-selectin blockade | Anti-mouse E-selectin (CD62E) | 9A9 | Rat IgG2b | Bio X Cell | BE0294 | AB_2687816 |
| E-selectin and ICAM-1 blockade control | Anti-keyhole limpet hemocyanin isotype control | LTF-2 | Rat IgG2b, $\kappa$ | Bio X Cell | BE0090 | AB_1107780 |
| ICAM-1 blockade | Anti-mouse ICAM-1 (CD54) | YN1/1.7.4 | Rat IgG2b, $\kappa$ | Bio X Cell | BE0020-1 | AB_1107661 |
| PECAM-1 blockade | Anti-mouse PECAM-1 (CD31) | 390 | Rat IgG2a, $\kappa$ | Bio X Cell | BE0377 | AB_2927514 |
| PECAM-1 blockade control | Anti-trinitroph enol isotype control | 2A3 | Rat IgG2a, $\kappa$ | Bio X Cell | BE0089 | AB_1107769 |
| P-selectin blockade | Anti-mouse/rat P-selectin (CD62P), Ultra-LEAF purified | RMP-1 | Mouse IgG2a, κ | BioLegend | 148308 | **AB_2565817** |
| P-selectin blockade control | Mouse IgG2a, κ isotype control, Ultra-LEAF purified | MOPC-173 | Mouse IgG2a, κ | BioLegend | 400264 | AB_11148947 |
| VCAM-1 blockade | Anti-mouse VCAM-1 (CD106) | M/K-2.7 | Rat IgG1, κ | Bio X Cell | BE0027 | AB_1107572 |
| VCAM-1 blockade control | Anti-horseradish peroxidase isotype control | HRPN | Rat IgG1, κ | Bio X Cell | BE0088 | AB_1107775 |
| F4/80 immunohistochemistry | Anti-mouse F4/80 | CI:A3-1 | Rat IgG2b | Novus Biologicals | NB600-404 | AB_10003219 |
| F4/80 staining control | Rat IgG2b isotype control | RTK4530 | Rat IgG2b, κ | BioLegend | 400602 | AB_326546 |
| F4/80 immunohistochemistry | Biotinylated goat anti-rat IgG (H+L), mouse adsorbed | Polyclonal | Goat | Vector Laboratories | BA-9401 | AB_2336208 |
| Intraepidermal nerve fibre staining | Anti-PGP9.5 | Polyclonal | Rabbit IgG | Dako/Agilent | Z5116 | AB_2622233 |
| Intraepidermal nerve fibre staining | Alexa Fluor 555 goat anti-rabbit IgG (H+L), cross-adsorbed | Polyclonal | Goat | Invitrogen/Thermo Fisher | A-21428 | AB_2535849 |

## Acknowledgements

We thank the IMB Advanced Microscopy Facility (IMB AMF) and the IMB Sequencing facility for technical support. We thank Ms. Andree Axelsen and the NHMRC (1186835) for the financial support of this study. Starobova was supported by NHMRC Investigator grant 2024/GNT2034754; Tay was supported by the Childhood Cancer Research Scholarship and the E.M.A and M.C Henker Postgraduate Medical Research Scholarship, Alshammari was supported by the University of Hafr Al Batin, Saudi Arabia scholarship; Meregalli was supported by EU in NextGenerationEU plan through the Italian “Bando Prin 2022D.D. 1409del 14-09-2022”. Shatunova was supported by the University of Queensland International Scholarship and Mater Foundation; Winkler was supported by Mater Research Institute, Mater Foundation, and Leukaemia Foundation, Australia and prior NHMRC;NIH grant 1130273;NS111929; Vetter was supported by NHMRC Investigator grant 2017086; Stow was supported by NHMRC APP1176209; Labzin was supported by an NHMRC Grant 2010757 and by the Australian Research Council (Future Fellowship FT240100816). This work was also supported by the Kids’ Cancer Project.

## Authors contributions

H.S., I.G.W. and I.V. conceived and supervised the study. H.S., T.J.P., and I.V. secured funding. H.S., A.A., N.T., S.S., A.L., F.I., V.R.-M., B.H., S.K. and D.L.B. performed experiments. A.A., H.S., Q.N., M.M.M., D.T-F., T.J.P. and N.N.I. performed data analysis, including spatial transcriptomic analyses. H.S., L.L., G.C., C.M., A.R., J.L.S., I.G.W. and I.V. provided key reagents, materials and resources. H.S. and I.V. wrote the manuscript with input from all authors. All authors reviewed and approved the final manuscript.

## Data availability

The spatial transcriptomics datasets generated in this study will be deposited in the Gene Expression Omnibus (GEO) prior to publication and will be made publicly available upon acceptance. Source data underlying all figures are provided with the manuscript in Supplementary tables. Additional data supporting the findings of this study are available from the corresponding authors upon reasonable request.

## Code availability

All analyses were performed using publicly available software, including DESeq2, SPOTlight, and CellChat. Custom scripts used for data processing, pseudobulk generation, and downstream analyses are available from the corresponding authors upon reasonable request.

