## Supplementary Materials for "E-Selectin Orchestrates IL-1β–Dependent Neuroinflammation via NLRP3 in Vincristine-Induced Neuropathy"

### 1. Supplementary Materials and methods

#### 1.1. Animal and ethical approvals

Behavioural and in vivo experiments were performed using 8–10-week-old C57BL/6J, E-selectin knockout (*Sele*$^{-/-}$), fucosyltransferase 4/7 double knockout (*Fut4/7*$^{-/-}$), and ASC–citrine reporter mice. Mice were group-housed (3–5 per cage) under a 12 h light/dark cycle with ad libitum food and water. All procedures were approved by the University of Queensland Animal Ethics Committee and conducted in accordance with the Animal Care and Protection Regulation (Qld, 2012), the Australian Code for the Care and Use of Animals for Scientific Purposes (2013), and IASP guidelines.

#### 1.2. Isolation and culture of bone Marrow-derived macrophages (BMDMs)

Murine (C57BL6/j or ASC-citrine) bone marrow-derived macrophages (BMDMs) were isolated and cultured as previously described[1,2]. Briefly, bone marrow was collected from 8-week-old male mice and differentiated over 7 days in RPMI-1640 medium (Gibco) supplemented with 10% heat-inactivated foetal bovine serum (FBS), 1% GlutaMAX (Life Technologies), and recombinant human CSF-1 (150 pg/mL). On day 7, cells were dissociated and seeded at a density of $1\times10^6$ cells/mL in Opti-MEM (Gibco) containing recombinant human CSF-1 (150 pg/mL) into 96-well plates. Plates were pre-coated for 24 h with one of the following: recombinant mouse E-selectin (5 µg/mL; R&D Systems), recombinant mouse P-selectin (5 µg/mL; R&D Systems), recombinant human IgG isotype control (5 µg/mL; Invitrogen), or Tris-buffered saline (PBS) as a control. After 1 h of cell adhesion, cells were primed with ultrapure lipopolysaccharide (LPS; 10 ng/mL; *E. coli* K12, tlrl-peklps; InvivoGen) for 3 h, pretreated with MCC950 (10 µM) or PBS for 30 min, followed by vincristine (100 µM; 6 h incubation, Pfizer), doxorubicin (50 µM; 6 h incubation; Sigma Aldrich) or cyclophosphamide (100 µM; 6 h incubation, Sigma Aldrich) or nigericin (5 µM; 45 min incubation; Sigma-Aldrich). Where indicated, the NLRP3 inhibitor MCC950 (10 µM)[3] was added 30 min prior to vincristine or nigericin treatment. Supernatants were collected for downstream cytokine analysis.

#### 1.3. Quantification of cytokine release from BMDMs

Cytokine and chemokine release was quantified by LEGENDplex™ Mouse Macrophage/Microglia Panel (BioLegend) according to the manufacturer's protocol. Readouts from samples were obtained using a flow cytometer (Beckman Coulter CytoFLEX-S). Data were analysed using the Data Analysis Software Suite by BioLegend. The concentration of interleukin-1 beta (IL-1β) from ASC-Citrine reporter BMDMs was quantified using or ELISA kit (R&D Systems) according to the manufacturer's protocol.

#### 1.4. Quantification of DRG neuron adhesion to E-selectin

Dorsal root ganglia (DRGs) or bone marrow-derived were isolated from 8-week-old male C57BL/6 mice as previously described [2,4-6]. Dissociated DRG neurons or BMDMs were seeded into 96-well plates pre-coated with recombinant mouse E-selectin, human IgG isotype control, or Tris-buffered saline (TBS) (5 µg/mL; 24 h at 4°C). Cell suspensions were added and incubated at for 1 h to allow adhesion. Non-adherent cells were removed by washing three times with fresh DMEM-high glucose (40 µL/well). Adherent cells were counted under a light microscope by a blinded observer.

#### 1.5. Quantification of ASC speck formation

BMDMs were isolated from ASC–citrine transgenic mice and differentiated for 7 days as described above. Live-cell imaging was performed every 10 min for up to 220 min using a Nikon Ti-E inverted microscope with a Hamamatsu Flash 4.0 sCMOS camera to monitor ASC speck formation. Supernatants were collected at the end and analysed for IL-1β by ELISA.

#### 1.6. Nf-κB p65 Transcription Factor Assay

NF-κB activation was assessed using the NF-κB p65 Transcription Factor Assay Kit (Abcam) following the manufacturer's instructions. Nuclear proteins were extracted 30 min post-treatment using the Abcam Nuclear Extraction Kit. Protein concentrations were measured via Nanodrop, and 20 µL of nuclear extract per sample was analysed in duplicate. Activated NF-κB p65 binding to DNA was detected using a specific antibody, HRP-conjugated secondary antibody, and a colorimetric readout at 450 nm. Absorbance was measured using a Tecan microplate reader, and data were analysed to compare NF-κB activity between conditions.

#### 1.7. Compounds and *in vivo* drug administration

Vincristine sulphate (Pfizer) was diluted in sterile phosphate-buffered saline (PBS) and administered intraperitoneally (i.p.) at 0.5 mg/kg (10 µl/g), as previously described[1]. To evaluate the role of cell adhesion molecules in VIPN, mice received two i.p. injections (24 h apart) of blocking antibodies against E-selectin, VCAM-1, ICAM-1, PECAM-1 (Bio X Cell), or P-selectin (BioLegend), or matching isotype controls (10 mg/kg; 10 µl/g), prior to vincristine administration. For IL-1 receptor inhibition, anakinra (100 mg/kg, Sobi) or PBS was diluted in sterile saline and administered i.p. at 24 h and 1 h before intraplantar (i.pl.) injection of recombinant mouse E-selectin. The specificity of all the blocking antibodies for their respective targets was validated *in vitro* using the BMDM adhesion assay (as above) prior to their use in in vivo experiments (data not shown). Recombinant mouse E-selectin (600 ng/injection; R&D Systems) or human IgG isotype control (600 ng/injection; Invitrogen) were diluted in sterile PBS and injected intraplantar (i.pl.; 20 µl/paw) into the right hind paw of C57BL6/J or *Fut4/7*$^{-/-}$ mice under 2% isoflurane anaesthesia.

#### 1.8. Intraplantar injection of bone Marrow-derived macrophages (BMDMs)

Bone marrow-derived macrophages (BMDMs) were differentiated from C57BL/6 mice in RPMI-1640 supplemented with CSF-1 for 7 days as described above. On day 7, BMDMs were seeded onto petri dishes pre-coated with recombinant mouse E-selectin or human IgG1 (both 5 µg/mL) for 1 h and stimulated as described above. After stimulation, cells were washed and resuspended in PBS. Prepared BMDMs (1,000 BMDMs in 20 µL/paw) were injected intraplantar (i.pl.) into C57BL6/J recipient mice under 2% isoflurane as described previously[1].

#### 1.9. Depletion of monocytes/macrophages

To deplete phagocytic cells, including monocytes and macrophages, mice received intraperitoneal (i.p.) injections of liposome-encapsulated clodronate (10 µl/g of 5 mg/mL; Liposoma) or control liposome-encapsulated PBS as described previously[1]. A second i.p. injection was administered 72 h later, concurrently with an intraplantar (i.pl.) injection of recombinant mouse E-selectin (600 ng in 20 µl; R&D Systems) into the hind paw. Spleens were collected post-treatment for histological analysis to confirm F4/80$^{+}$ cell depletion.

#### 1.10. Immunohistochemistry staining

Dorsal root ganglia and sciatic nerves were collected 25 h post vincristine or vehicle injection, fixed in 10% neutral-buffered formalin (NBF), and embedded in paraffin as previously described[7]. Briefly, Paraffin sections (5 µm) were deparaffinised, rehydrated, and antigen-retrieved with Proteinase K. After blocking with Background Sniper (BioCare Medical), sections were incubated with rat anti-mouse F4/80 (1:350; Novus Bio) for 2 h, followed by biotinylated secondary antibody, ABC amplification (Agilent Dako), and DAB detection. Nuclei were counterstained with haematoxylin. Isotype control (rat IgG2b; BioLegend) was used to confirm specificity. Slides were scanned (Olympus) and F4/80+ area quantified using Visiopharm by a blinded observer to assess macrophage infiltration across treatment groups.

#### 1.11. Mechanical and thermal paw withdrawal threshold measurements

Mechanical sensitivity of the hind paws was assessed using an electronic von Frey apparatus (MouseMet; Topcat Metrology) as previously described[8]. Mice were acclimatised in MouseMet enclosures (bar-bottom floor) for 30 min. Only responses within preset linear parameters were included. PWT was recorded automatically as withdrawal force (g). Each biological replicate represented the average of three measurements per mouse, taken ≥5 min apart. Thermal sensitivity of then hind paws was assessed using the MouseMet Thermal device (Topcat Metrology, UK) as previously described[9]. The probe, preheated to ~37°C, was applied to the plantar hind paw, triggering a controlled heat ramp (2.5°C/sec) upon light contact (~1 g force). Heating stopped automatically at paw withdrawal or when the probe was removed, with the withdrawal temperature (PWT) displayed digitally. A cut-off of 60°C was set to prevent tissue damage. Each biological replicate represented the average of three measurements per mouse, taken ≥5 min apart.

#### 1.12. Tissue collection for spatial transcriptomics and RNA quality control

Eight-week-old male C57BL/6J mice were treated with either saline or vincristine (0.5 mg/kg, i.p.) once. Additional groups included C57BL6/J mice pre-treated with an E-selectin–blocking antibody (Bio X Cell, 200 µg/mouse, i.p. at two time points (-24h ,-1h) prior to vincristine), and E-selectin knockout ($Sele^{-/-}$) mice treated once with vincristine (0.5 mg/kg, i.p.). Dorsal root ganglia (DRG) were collected 24h post vincristine treatment under RNase-free conditions and immediately embedded in chilled OCT compound (Tissue-Tek, Sakura, Japan), then flash-frozen in pre-cooled isopentane to preserve tissue morphology. The study included DRGs from lumbar level L3-L5 (both sides) from 3 mice per treatment condition. A total of 15 DRGs per condition were embedded in one OCT block, positioned on a uniform embedding plane for consistent sectioning. Blocks were stored at –80 °C until further processing.

Prior to spatial transcriptomics, RNA quality and optimal sectioning parameters were assessed. RNA was extracted from 30–40 µm scrolls using the RNA Micro Kit (Qiagen), and RNA integrity was confirmed with an Agilent 6000 Pico Kit on a Bioanalyzer; all samples had RIN > 7.0. Sectioning and permeabilization times were optimized using Visium Tissue Optimization Slides (10x Genomics, User Guide Rev E). Both 8 µm and 10 µm sections yielded robust cDNA signals with 28–30 min permeabilization; 10 µm sections were selected to approximate single-cell thickness.

#### 1.13. Library preparation and sequencing

Cryosections were dried at 37 °C for 1 min, fixed in 100% methanol at –20 °C for 30 min, and stained with Mayer's haematoxylin (5 min) and eosin (2 min). Slides were mounted in 85% glycerol for imaging on a Zeiss Axio Z1 slide scanner. Library preparation followed the Visium Spatial Gene Expression User Guide (CG000239 Rev C, 10x Genomics). Sequencing was performed using an Illumina NextSeq 500 with the following read structure: Read1 – 28 bp,

Index1 – 10 bp, Index2 – 10 bp, Read2 – 120 bp. Raw BCL files were processed using bcl2fastq (v2.7.0), and reads were aligned to the mouse reference genome using Space Ranger (v1.3.0, 10x Genomics).

#### 1.14. Quantification of Intraepidermal Nerve Fibre (IENF) density

Vincristine (0.5 mg.kg. i.p.; Pfizer) was injected 5 times a week for 4 weeks. 1 week after the final vincristine injection plantar hind paw skin was collected, postfixed in 4% paraformaldehyde for 24 h at 4 °C, cryoprotected in 30% sucrose o/n, embedded in OCT (TissueTEK) and snap frozen using dry ice[1]. Tissue was sectioned into three non-consecutive 50 µm sections per animal (n = 4 animals per group) and mounted onto poly-D-lysine–coated slides. Sections were permeabilized with 0.1% Triton X-100, quenched with ammonium chloride (50mM in PBS, 30 min at RT) and blocked with 0.5% BSA before incubation with rabbit anti-PGP9.5 primary antibody (1:500; Dako, Z5116) followed by Alexa Fluor 555–conjugated goat anti-rabbit secondary antibody (1:1000; Invitrogen, A-21428). Sections were coverslipped with DAPI-containing mounting media (Fluoroshield, Sigma Aldrich, F6057). Images were acquired using a spinning-disc confocal microscope (Nikon Ti2/Andor Dragonfly). Image analysis was performed in FIJI (ImageJ). Intraepidermal nerve fibre density was expressed as the percentage of PGP9.5-positive area relative to total epidermal area.

### 2. Supplementary Figures

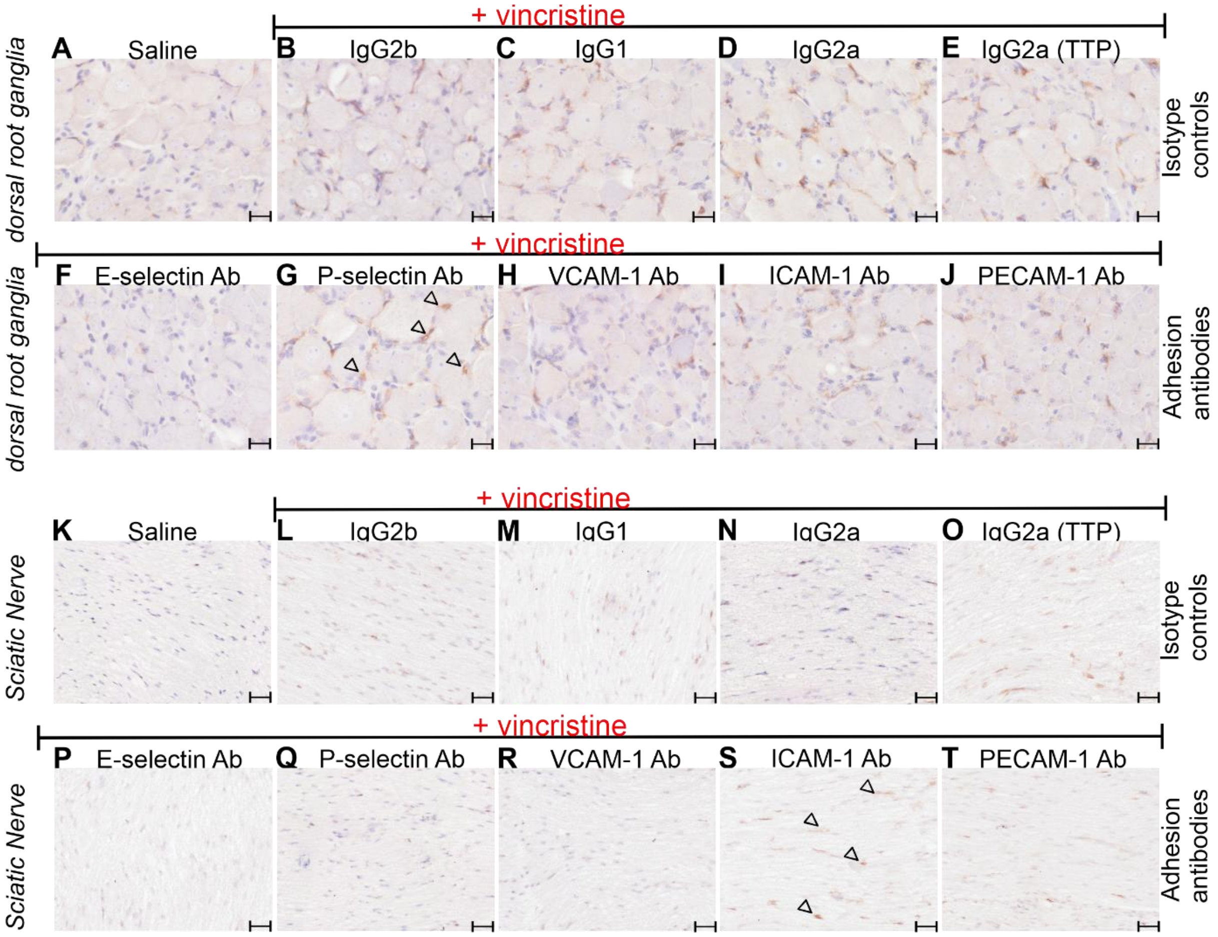


***Supplementary Fig. S1. Representative immunohistochemical images of F4/80⁺ area in dorsal root ganglia and sciatic nerve 25 h post-vincristine administration.*** *Mice received two intraperitoneal (i.p.) injections of antibodies targeting E-selectin (E-sel Ab), P-selectin*

*(P-sel Ab), VCAM-1 (VCAM-1 Ab), ICAM-1 (ICAM-1 Ab), and PECAM-1 (PECAM-1 Ab), or corresponding isotype control antibodies (IgG2B, IgG1, IgG2A(TTP)) (10 mg/kg; i.p.) at two time points (t -24h, t -1h). This was followed by a single vincristine injection (0.5 mg/kg, i.p., time point t 0h). Shown are representative pictures of F4/80$^+$ area in dorsal root ganglia (**A-J,** top panel: isotype controls; second panel: selective adhesion antibodies) and sciatic nerve (**K-T**) in a saline control group (**A**; **K**) and following pre-treatment with isotype control antibodies (**B-E**; **L-O**) or adhesion molecule-inhibiting antibodies (**F-J**;**P-T**). Scale bar = 20 µm for DRGs or 10 µm for SN. Examples of F4/80$^+$ area (brown stain) indicated by arrowheads.*

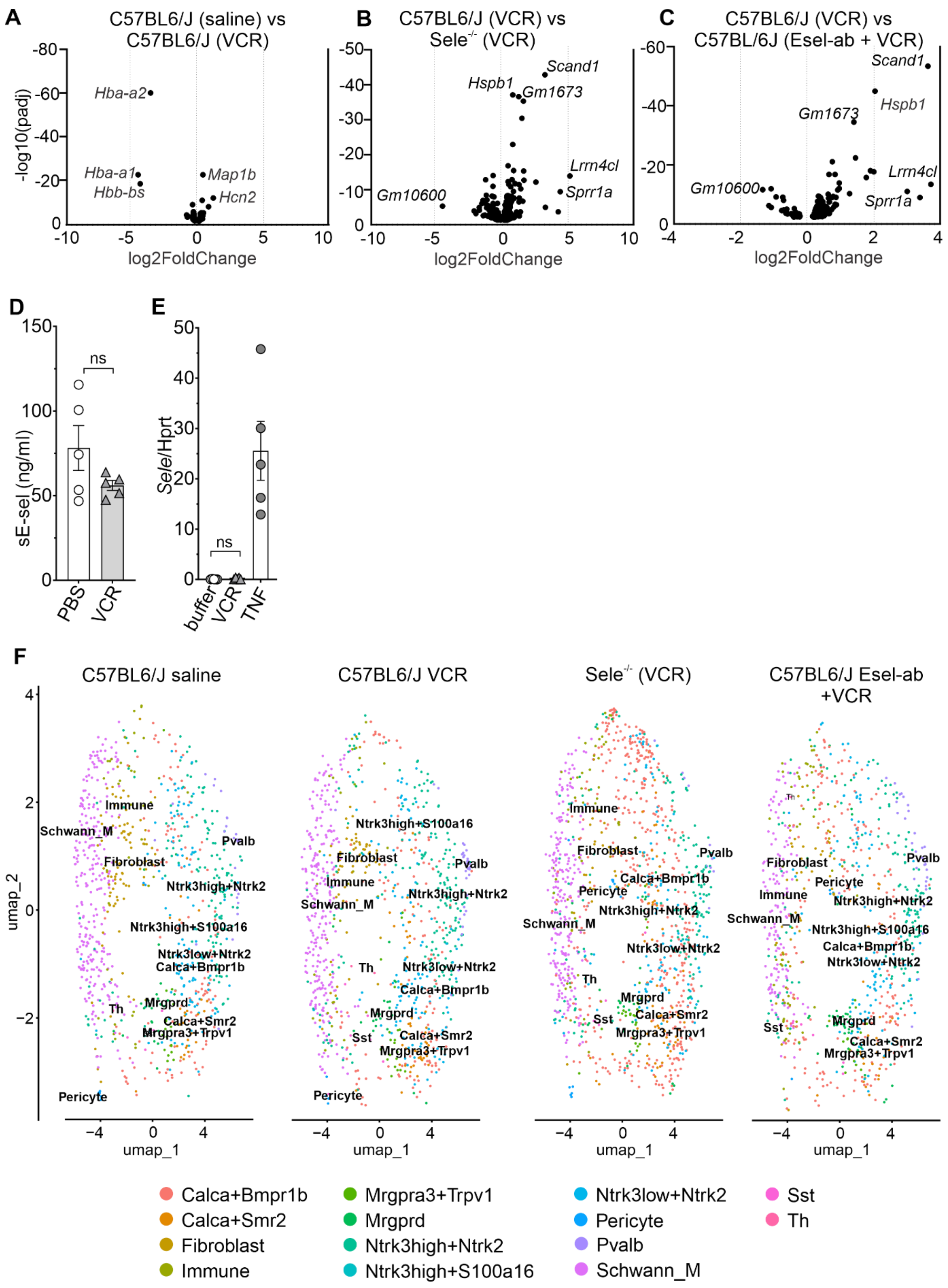


***Supplementary Fig. S2. Vincristine induces a stress and injury transcriptional programme in dorsal root ganglia without upregulating E-selectin. (A–C)*** *Volcano plots of pseudobulk differential gene expression in dorsal root ganglia (DRG).* ***(A)*** *C57BL6/J saline versus vincristine (VCR) reveals induction of injury- and excitability-associated genes (e.g. Map1b, Hcn2) and downregulation of haemoglobin genes (Hba-a1, Hba-a2, Hbb-bs).* ***(B)*** *C57BL6/J*

*(VCR) versus* $Sele^{-/-}$*(VCR) and* ***(C)*** *C57BL6/J (VCR) versus C57BL6/J E-selectin antibody pre-treatment (Esel-ab + VCR) identify stress-related transcripts (e.g. Hspb1, Scand1) associated with E-selectin perturbation.* ***(D–E)*** *Circulating soluble E-selectin plasma protein levels in C57BL6/J mice treated with vincristine (0.5 mg.kg, i.p., once) and Sele/Hprt transcript expression in HUVEC cells treated with vincristine for 24 hours, show no induction of Sele expression by vincristine, while Tnf serves as a positive control (ns, not significant).* ***(F)*** *UMAP projections of cell types across conditions (saline, VCR,* $Sele^{-/-}$ *VCR, Esel-ab + VCR) demonstrate preservation of major neuronal and non-neuronal cell populations. Abbreviations: VCR: vincristine;* $Sele^{-/-}$*: E-selectin–deficient mice; Esel-ab: E-selectin–blocking antibody; DRG: dorsal root ganglion; TNF: tumour necrosis factor; Schwann_M: myelinating Schwann cells;* $Calca^{+}$*Bmpr1b and* $Calca^{+}$*Smr2: peptidergic sensory neuron subtypes;* $Mrgpra3^{+}$*Trpv1: nociceptive/pruriceptive neurons; Mrgprd: non-peptidergic sensory neurons; Ntrk3^high/+Ntrk2 and Ntrk3^low/+Ntrk2: sensory neuron subsets; Pvalb: parvalbumin-expressing neurons; Sst: somatostatin-expressing neurons; Th: tyrosine hydroxylase–expressing neurons; Immune: aggregated immune cell populations; sE-sel: soluble E-selectin.*

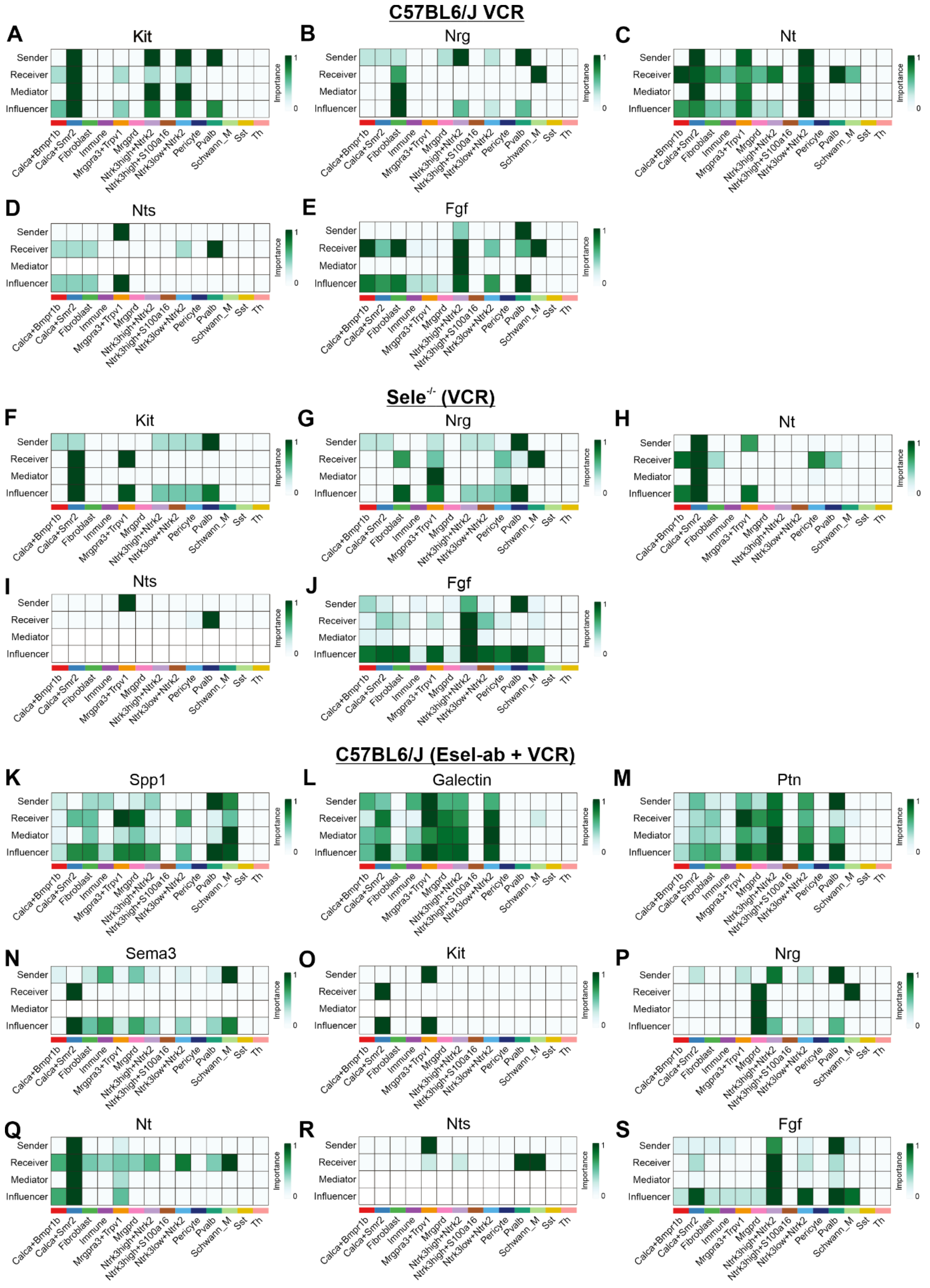


***Supplementary Fig. S3: Vincristine and E-selectin perturbation reshape growth factor and neuromodulatory communication networks in dorsal root ganglia.*** *CellChat analysis was used to infer ligand–receptor–mediated communication networks in dorsal root ganglia (DRG) following vincristine treatment, highlighting pathway-specific sender, receiver, mediator and*

*influencer roles across neuronal and non-neuronal cell populations.* ***(A–E)*** *Growth factor and neuromodulatory signalling pathways in C57BL6/J mice treated with vincristine (VCR). Heatmaps depict the relative contribution of each annotated cell type as senders, receivers, mediators or influencers for KIT (A), NRG (B), NT (C), NTS (D) and FGF (E) pathways, with darker green indicating higher importance scores.* ***(F–J)*** *Corresponding analyses in* $Sele^{-/-}$ *mice treated with vincristine (VCR) show preservation of KIT, NRG, NT, NTS and FGF pathways but redistribution of communication roles across neuronal, glial and stromal populations.* ***(K–S)*** *Analyses in C57BL6/J mice treated with vincristine following E-selectin antibody pre-treatment (Esel-ab + VCR). Heatmaps illustrate altered sender, receiver, mediator and influencer contributions for SPP1 (K), GALECTIN (L), PTN (M), SEMA3 (N), KIT (O), NRG (P), NT (Q), NTS (R) and FGF (S) pathways, highlighting further redistribution of signalling roles relative to genetic E-selectin deletion. Across all panels, rows indicate inferred communication roles (sender, receiver, mediator, influencer) and columns represent annotated DRG cell populations; colour intensity reflects relative importance within each pathway. VCR: vincristine;* $Sele^{-/-}$*: E-selectin–deficient mice; DRG: dorsal root ganglion; Schwann_M: myelinating Schwann cells;* $Calca^{+}$*Bmpr1b and* $Calca^{+}$*Smr2: peptidergic sensory neuron subtypes expressing Calca with Bmpr1b or Smr2, respectively; Mrgprd: non-peptidergic sensory neurons; Mrgpra3*$^{+}$*Trpv1: nociceptive/pruriceptive neurons; Ntrk3^high/+Ntrk2 and Ntrk3^low/+Ntrk2: sensory neuron subsets defined by relative Ntrk3 and Ntrk2 expression; Pvalb: parvalbumin-expressing neurons; Sst: somatostatin-expressing neurons; Th: tyrosine hydroxylase–expressing neurons; Immune: aggregated immune cell populations; KIT: stem cell factor receptor signalling; NRG: neuregulin signalling; NT: neurotrophin signalling; NTS: neurotensin signalling; FGF: fibroblast growth factor signalling; SPP1: osteopontin signalling; PTN: pleiotrophin signalling; SEMA3: semaphorin 3 signalling.*

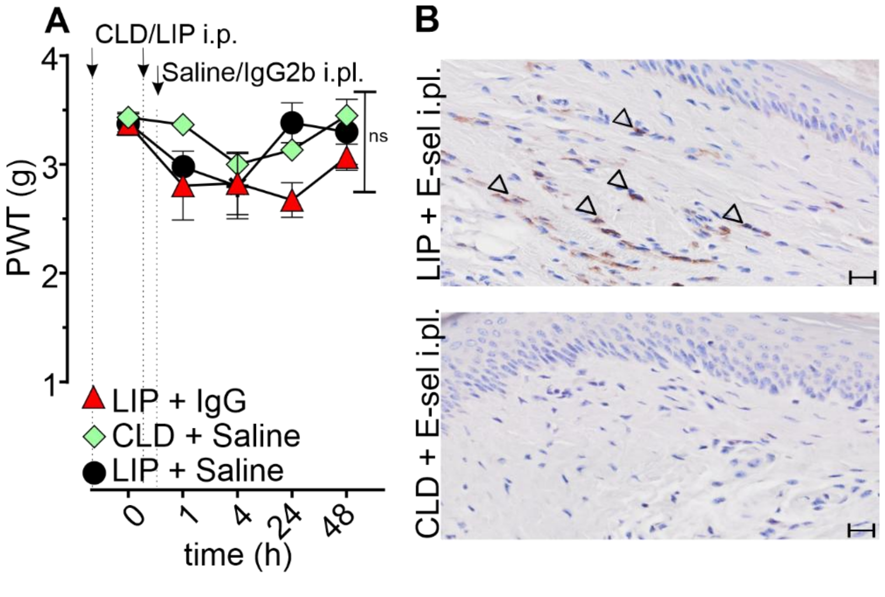


***Supplementary Fig. S4.:*** ***(A)*** *Control groups treated with liposomes (LIP), clodronate liposomes (CLD), saline or IgG2b did not develop mechanical hypersensitivity.* ***(B)*** *Representative images of* $F4/80^{+}$ *immunohistochemistry in footpad skin of mice pretreated with CLD or LIP and injected with i.pl. E-selectin. Statistical significance (#: P < 0.05) was determined using repeated measures two-way ANOVA (n = 6-9).*

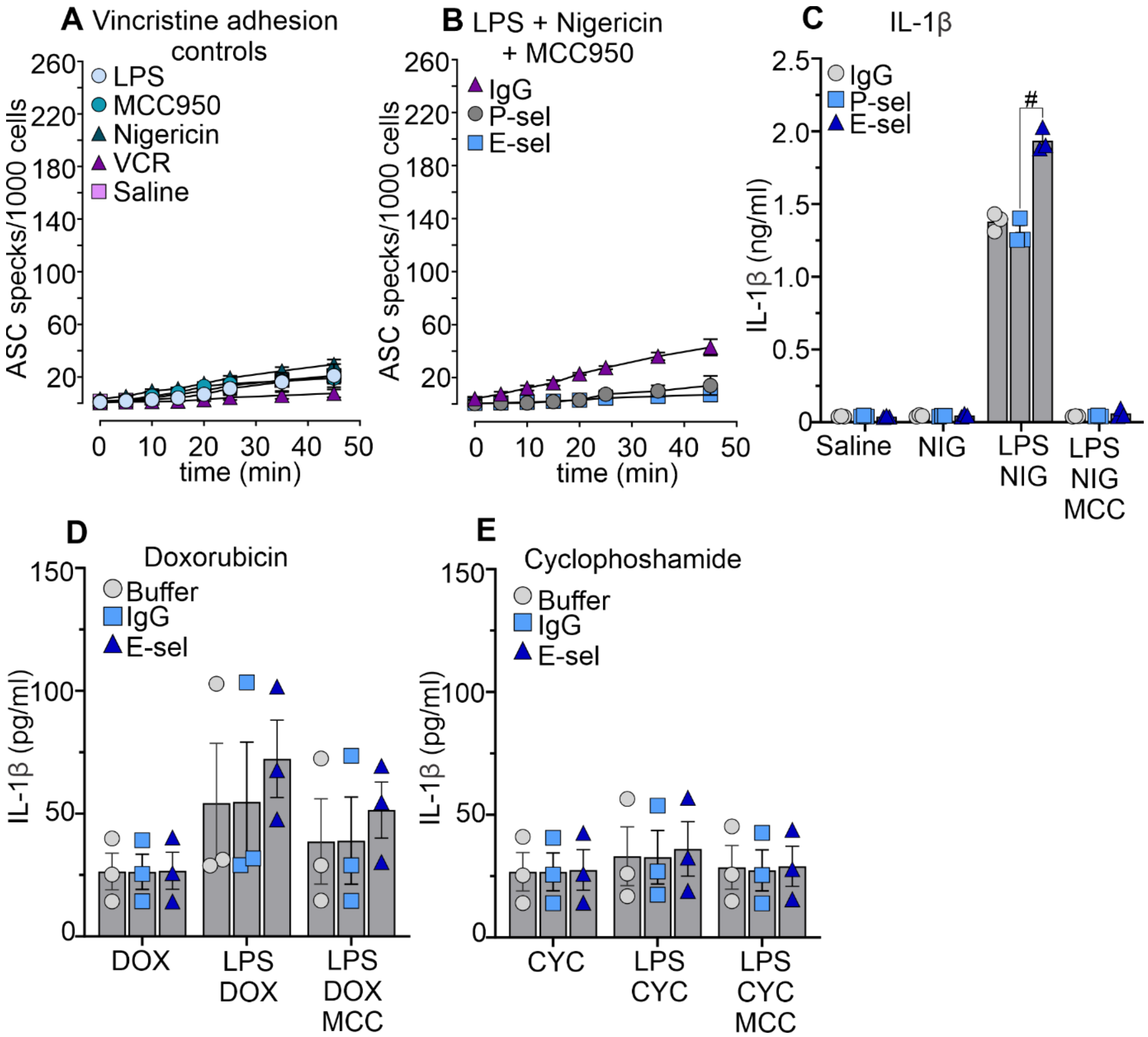


***Supplementary Figure S5. E-selectin enhances ASC speck formation and IL-1β release in vincristine- or nigericin-treated BMDMs.*** *BMDMs were adhered to E-selectin-, P-selectin- or IgG-coated surfaces (5 µg/mL), primed with LPS (10 ng/mL, 3 h), and treated as indicated. MCC950 (10 µM) was added 30 min before stimulation; VCR (100 µM), doxorubicin (DOX; 50 µM) and cyclophosphamide (CYC; 100 µM) were applied for 6 h, and nigericin (5 µM) for 45 min. (**A**) No significant ASC speck formation was observed in E-selectin–adhered BMDMs treated with LPS, MCC950, nigericin, vincristine (VCR), or saline alone. (**B,C**) The NLRP3 inhibitor MCC950 (10 µM) prevents ASC speck formation (**B**) and IL-1β release (**C**) induced by treatment with lipopolysaccharide (LPS, 10ng/ml) + nigericin (5 µM). (**D–E**) IL-1β release from BMDMs treated with LPS + doxorubicin (DOX, 50 µM) (D) or LPS + cyclophosphamide (CYC, 100 µM) (E) adhered to E-sel, IgG2b, or P-sel. No significant differences were observed across conditions. Statistical analysis: two-way ANOVA (**A-B**), one-way (**C,D,E**) (#P < 0.05); n≥3.*
