## Supplementary Figures for "E-Selectin Orchestrates IL-1β–Dependent Neuroinflammation via NLRP3 in Vincristine-Induced Neuropathy": Supplementary Figures.pdf

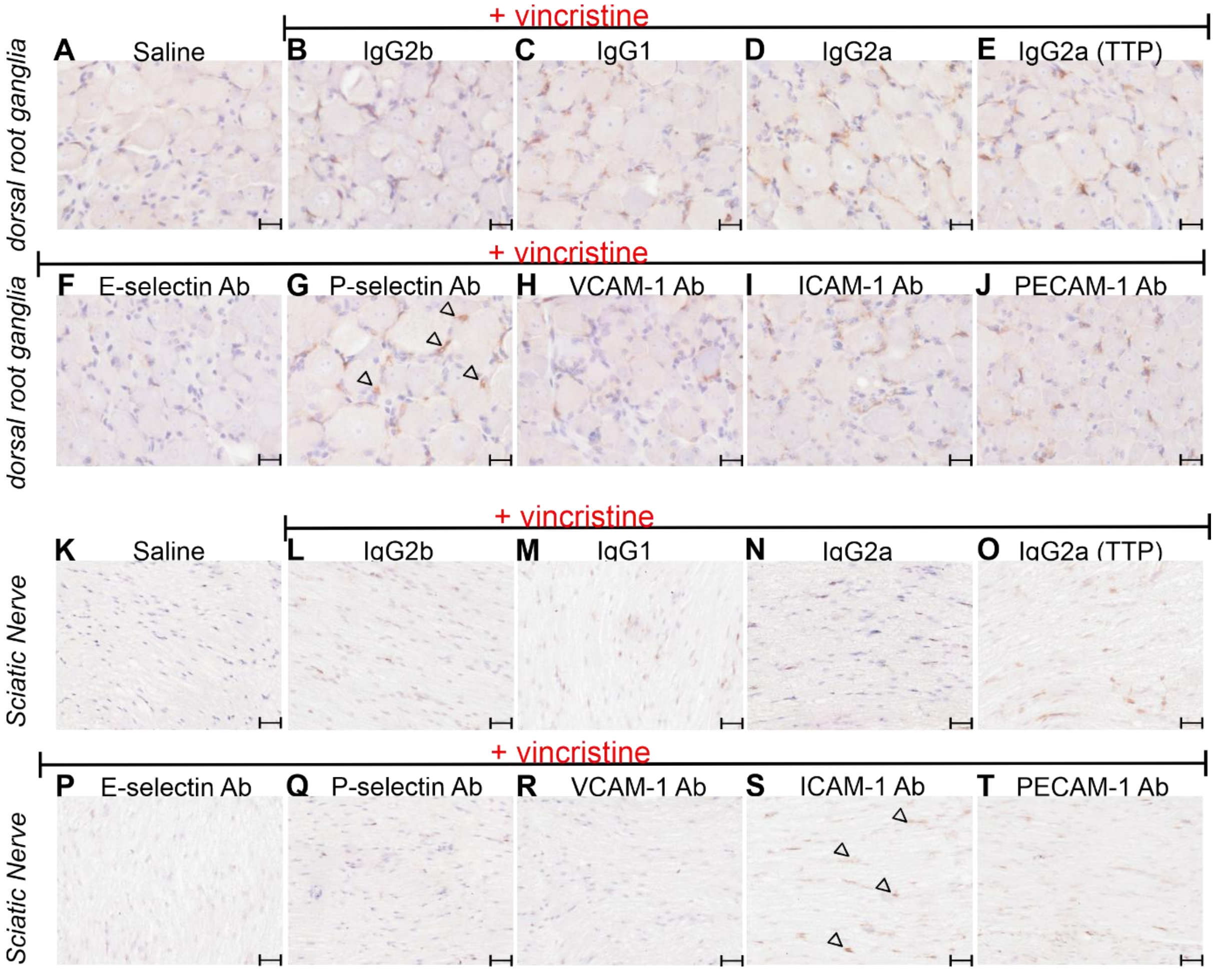


***Supplementary Fig. S1. Representative immunohistochemical images of F4/80⁺ area in dorsal root ganglia and sciatic nerve 25 h post-vincristine administration.*** *Mice received two intraperitoneal (i.p.) injections of antibodies targeting E-selectin (E-sel Ab), P-selectin (P-sel Ab), VCAM-1 (VCAM-1 Ab), ICAM-1 (ICAM-1 Ab), and PECAM-1 (PECAM-1 Ab), or corresponding isotype control antibodies (IgG2B, IgG1, IgG2A(TTP)) (10 mg/kg; i.p.) at two time points (t -24h, t -1h). This was followed by a single vincristine injection (0.5 mg/kg, i.p., time point t 0h). Shown are representative pictures of F4/80⁺ area in dorsal root ganglia (**A-J**) and sciatic nerve (**K-T**) in a saline control group (**A**; **K**) and following pretreatment with isotype control antibodies (**B-E**; **L-O**) or adhesion molecule-inhibiting antibodies (**F-J**;**P-T**). Scale bar = 20 µm for DRGs or 10 µm for SN. Examples of F4/80⁺ area (brown stain) indicated by arrowheads.*

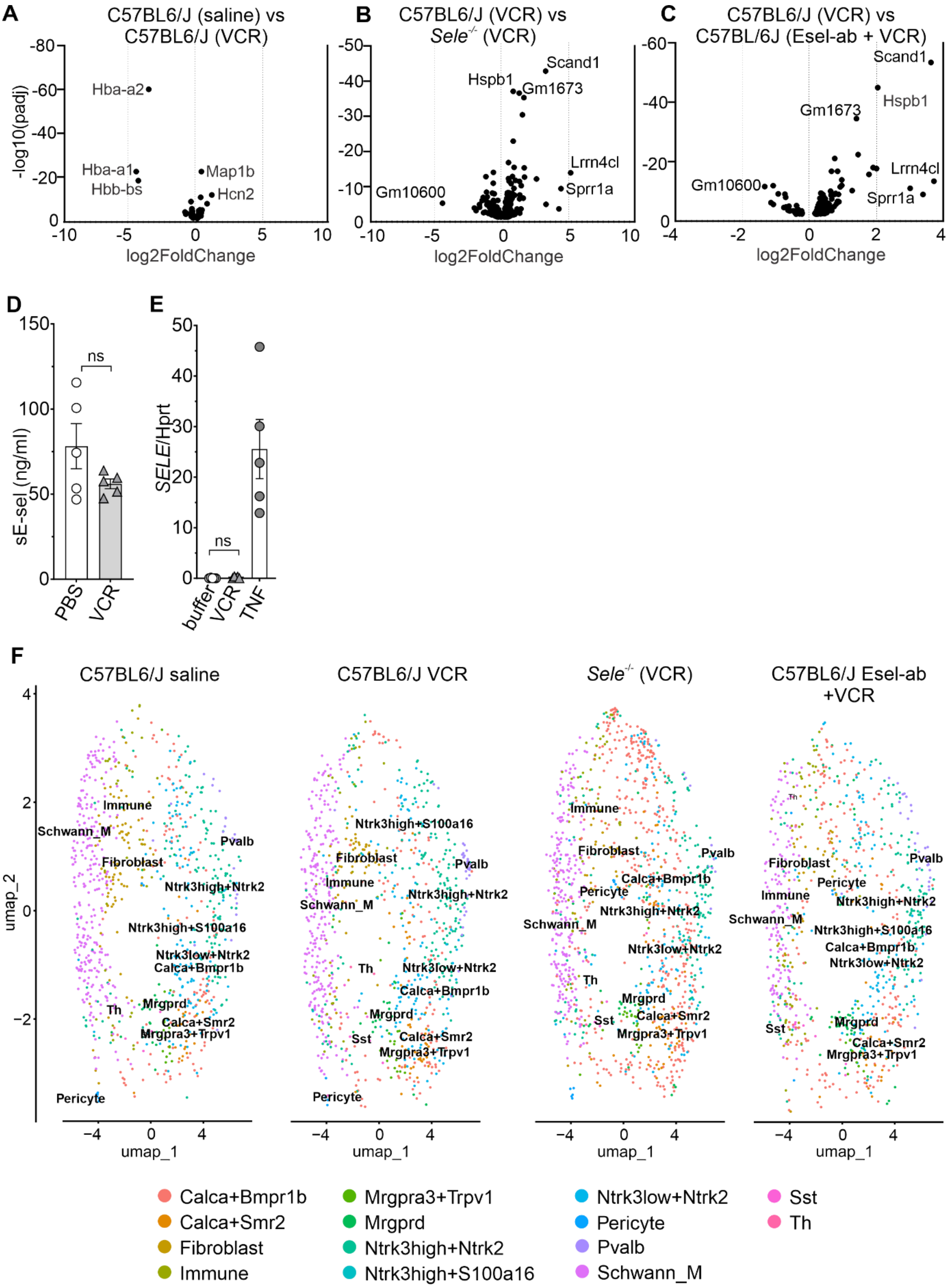


***Supplementary Fig. S2. Vincristine induces a stress and injury transcriptional programme in dorsal root ganglia without upregulating E-selectin. (A–C)*** *Volcano plots of pseudobulk differential gene expression in dorsal root ganglia (DRG).* ***(A)*** *C57BL6/J saline versus vincristine (VCR) reveals induction of injury- and excitability-associated genes (e.g. Map1b, Hcn2) and downregulation of haemoglobin genes (Hba-a1, Hba-a2, Hbb-bs).* ***(B)*** *C57BL6/J*

*(VCR) versus Sele$^{-/-}$ (VCR) and* ***(C)*** *C57BL6/J (VCR) versus C57BL6/J E-selectin antibody pre-treatment (Esel-ab + VCR) identify stress-related transcripts (e.g. Hspb1, Scand1) associated with E-selectin perturbation.* ***(D–E)*** *Circulating soluble E-selectin protein levels in C57BL6/J mice treated with vincristine (0.5 mg.kg, i.p., once) and SELE transcript expression in HUVEC cells treated with vincristine for 24 hours, show no induction of SELE expression by vincristine, while TNF serves as a positive control (ns, not significant).* ***(F)*** *UMAP projections of cell types across conditions (saline, VCR, Sele$^{-/-}$ VCR, Esel-ab + VCR) demonstrate preservation of major neuronal and non-neuronal cell populations. Abbreviations: VCR: vincristine; Sele$^{-/-}$: E-selectin–deficient mice; Esel-ab: E-selectin–blocking antibody; DRG: dorsal root ganglion; TNF: tumour necrosis factor; Schwann_M: myelinating Schwann cells; Calca$^{+}$Bmpr1b and Calca$^{+}$Smr2: peptidergic sensory neuron subtypes; Mrgpra3$^{+}$Trpv1: nociceptive/pruriceptive neurons; Mrgprd: non-peptidergic sensory neurons; Ntrk3^high/+Ntrk2 and Ntrk3^low/+Ntrk2: sensory neuron subsets; Pvalb: parvalbumin-expressing interneurons; Sst: somatostatin-expressing neurons; Th: tyrosine hydroxylase–expressing neurons; Immune: aggregated immune cell populations; sE-sel: soluble E-selectin.*

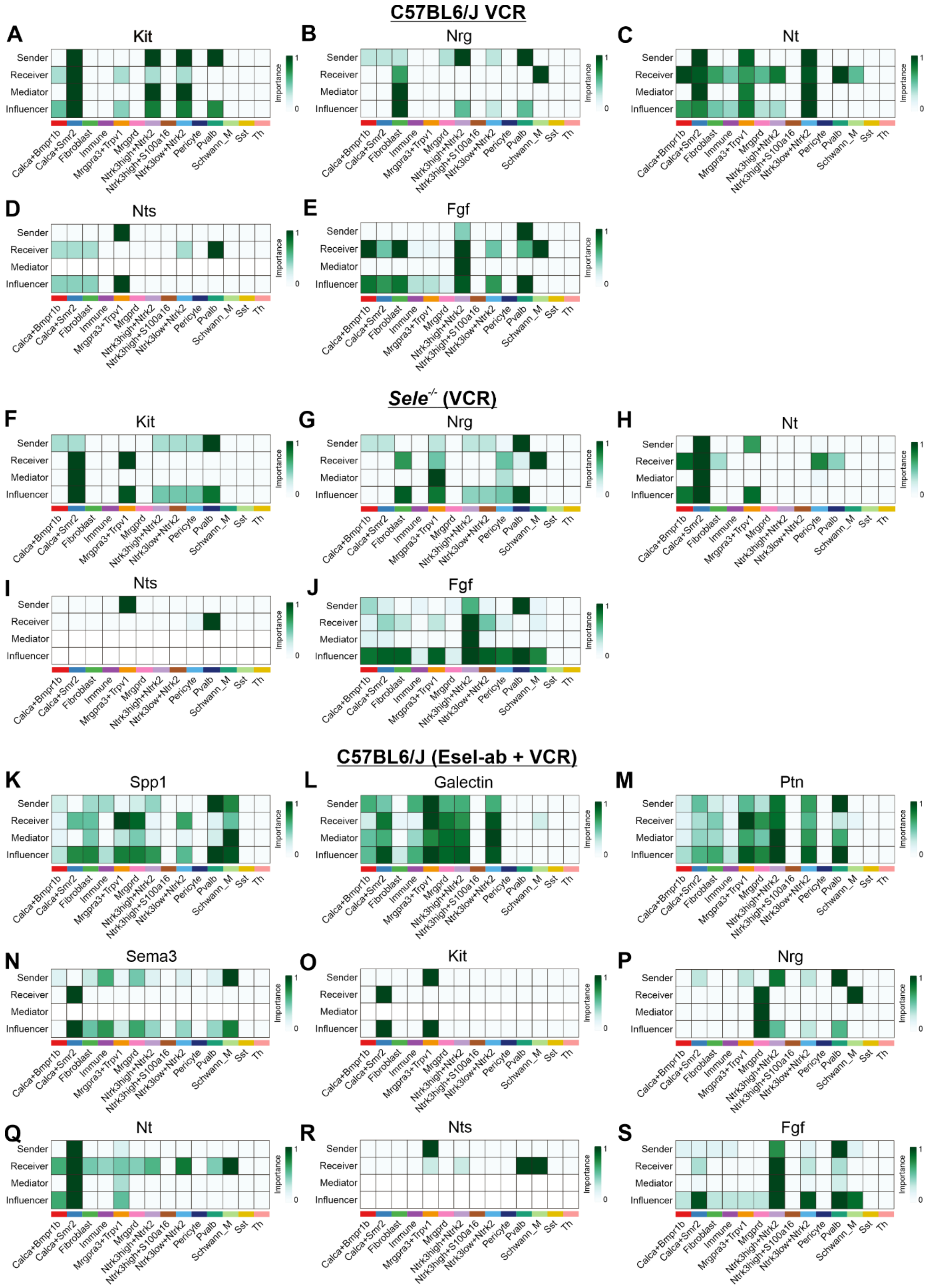


***Supplementary Fig. S3: Vincristine and E-selectin perturbation reshape growth factor and neuromodulatory communication networks in dorsal root ganglia.*** *CellChat analysis was used to infer ligand–receptor–mediated communication networks in dorsal root ganglia (DRG) following vincristine treatment, highlighting pathway-specific sender, receiver, mediator and influencer roles across neuronal and non-neuronal cell populations.* ***(A–E)*** *Growth factor and*

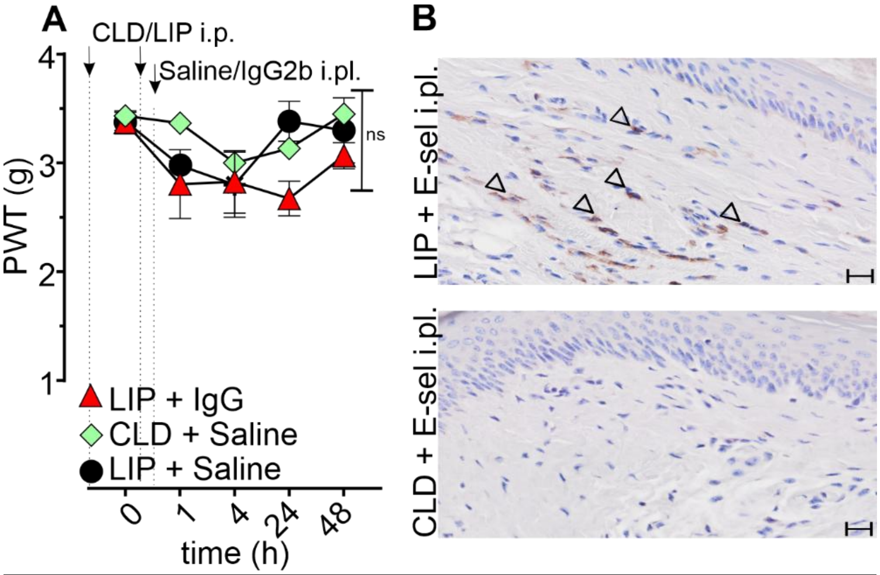


***Supplementary Fig. S4.:*** ***(A)*** *Control groups treated with liposomes (LIP), clodronate liposomes (CLD), saline or IgG2b did not develop mechanical hypersensitivity.* ***(B)*** *Representative images of $F4/80^{+}$ immunohistochemistry in footpad skin of mice pretreated with CLD or LIP and injected with i.pl. E-selectin. Statistical significance (#: $P < 0.05$) was determined using repeated measures two-way ANOVA (n = 6-9).*

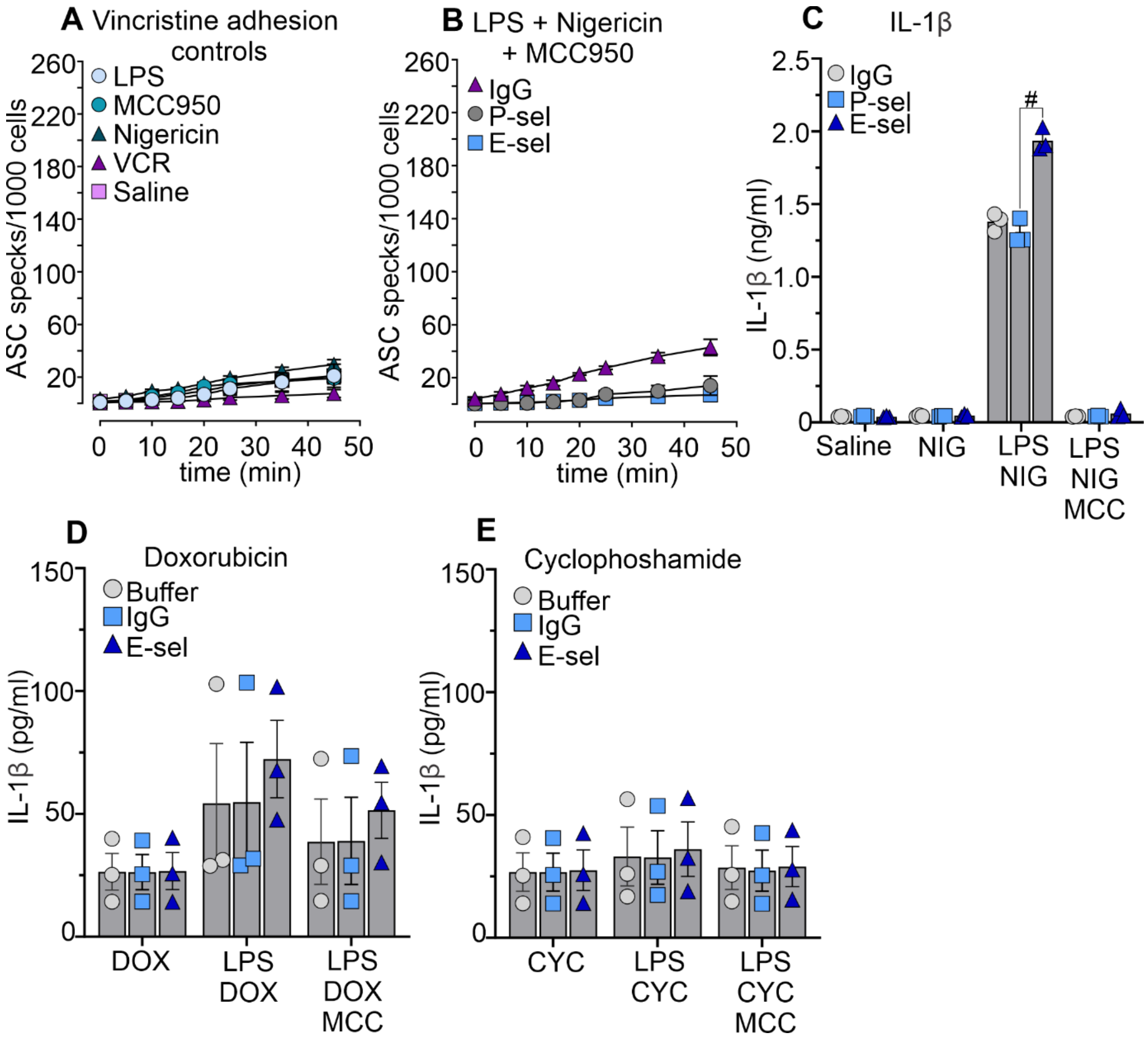


***Supplementary Figure S5. E-selectin enhances ASC speck formation and IL-1β release in vincristine- or nigericin-treated BMDMs.*** *(**A**) No significant ASC speck formation was observed in E-selectin–adhered BMDMs treated with LPS, MCC950, nigericin, vincristine (VCR), or saline alone. (**B,C**) The NLRP3 inhibitor MCC950 (10 µM) prevents ASC speck formation (**B**) and IL-1β release (**C**) induced by treatment with lipopolysaccharide (LPS, 10ng/ml) + nigericin (5 µM). (**D–E**) IL-1β release from BMDMs treated with LPS + doxorubicin (DOX, 50 µM) (D) or LPS + cyclophosphamide (CYC, 100 µM) (E) adhered to E-sel, IgG2b, or P-sel. No significant differences were observed across conditions. Statistical analysis: one-way (**C,D,E**) or two-way (A,B) ANOVA with Dunnett's test (#$P < 0.05$); $n \geq 6$.*

**Intraplantar injection of bone Marrow-derived macrophages (BMDMs)**

Bone marrow-derived macrophages (BMDMs) were differentiated from C57BL/6 mice in RPMI-1640 supplemented with CSF-1 for 7 days as described above. On day 7, BMDMs were seeded onto petri dishes pre-coated with recombinant mouse E-selectin or human IgG1 (both 5 μg/mL) for 1 h at 4°C and stimulated as described above. After stimulation, cells were washed and resuspended in PBS at ~40 000 cells/mL. Prepared BMDMs (25 μl/paw) were injected intraplantar (i.pl.) into C57BL/6 recipient mice under 2% isoflurane as described previously[8].

**Pseudo-bulk generation from Visium data and differential expression analysis**

To obtain biologically independent replicates, Visium barcodes belonging to the same animal were summed to create one pseudo-bulk count matrix per animal. Aggregating at the animal level minimizes within-animal sampling noise and ensures that statistical inference pertains to variation across animals rather than across individual barcodes. Raw gene counts were analyzed in DESeq2 (Love et al., 2014). Size factors were estimated with the median-ratio method (each gene count divided by a barcode-specific factor derived from the geometric mean across samples), after which a batch term was included in the design formula to correct for sequencing batch effects (design = ~ batch + condition). DESeq2 fits a negative binomial generalized linear model and applies Wald statistics followed by Benjamini-Hochberg correction to identify differentially expressed genes. $Log_2$ fold-changes were shrunken with "ashr" (Stephens, 2017), improving effect-size ranking for low-count genes. Genes meeting p-adjusted $P \leq 0.05$ and $log_2FC \geq 0.585$(FC of 1.5) were reported as differentially expressed.

**Spatial Transcriptomics Deconvolution with SPOTlight**

Seeded non-negative matrix factorization (NMF) regression-based deconvolution was performed using the **SPOTlight** package (1.8.0) (Elosua-Bayes et al., 2021). Cell type proportions were inferred in the spatial transcriptomics dataset using a reference single-cell RNA-seq dataset containing previously annotated cell populations (Bhuiyan et al., 2024). Gene expression and metadata were extracted from the spatial transcriptomics data to construct a SpatialExperiment object. SPOTlight was then used to generate a deconvolution matrix based on the marker gene profiles from the reference single-cell clusters. Topic profiles representative of each cell type was projected onto spatial transcriptomic spots to estimate cellular composition.

**Data and statistical analyses**

Data (except transcriptomic analyses) were analysed using GraphPad Prism v10. Statistical significance as indicated in figures was set at adjusted $P < 0.05$. Behavioural and time-course data were assessed using repeated-measures two-way ANOVA with Šidák's post hoc test. One-way ANOVA with Šidák's test was used for immunohistochemistry, ELISA, and LEGENDPlex cytokine assays. Unpaired *t*-tests were used for two-group comparisons. Data are presented as mean ± SEM, with a minimum of six animals per group ($n \geq 6$). Comparisons were made against vehicle- or wildtype controls as appropriate.
